# Roles of *spoVF* operon subunits A and B in regulating cell morphology and biofilm formation in the biocontrol agent *Bacillus subtilis*

**DOI:** 10.1101/2024.12.18.629248

**Authors:** Yang Yang, Yu-Xin Zhou, Le-Yao Jia, Xin-Yang Song, Ya-Jie Lai, Su-Yan Wang, Pedro Laborda, Xin-Chi Shi

## Abstract

Enhanced expression of *spoVF* operon (sporulation initiation factor; dipicolinic acid (DPA) synthase) in *Bacillus subtilis* vegetative cells has been reported to increase host attachment and biocontrol ability. However, how the *spoVF* operon regulates these essential biocontrol factors remains unknown. In this study, it was found that the knockout of the *spoVFA* or *spoVFB* subunits, individually (*Δ*A and *Δ*B), or together (*Δ*A*Δ*B), decreased DPA biosynthesis, host attachment, nutrient competition ability, and biocontrol efficacy in *B. subtilis*. Moreover, the cell shape of the knockout strains changed from short rod to spherical. qRT-PCR analysis indicated that the *spoVF* operon regulates TCA cycle expression and biofilm formation by balancing the amounts of various transcriptional factors, such as spo0A, SinR, and AbrB, and the cofactor NADPH. The regulatory mechanism of the *spoVF* operon in *B. subtilis* biofilm-related genes was proposed for the first time.

**IMPORTANCE:** *Bacillus subtilis SpoVF* operon contains the *spoVFA* or *spoVFB* subunits, which are responsible for the dipicolinic acid synthase activity. Dipicolinic acid is an important component of *B. subtilis* spores. In this study, the roles of *spoVFA* or *spoVFB* subunits in the biocontrol ability *B. subtilis* were examined, indicating that the *spoVFA* subunit plays a key role in *B. subtilis* biofilm formation, cell morphology, growth, and nutrient competition ability. Additionally, the regulatory network connecting the *spoVF* operon with biofilm formation-related genes of *B. subtilis* was proposed. The results obtained in this study help to understand key genes involved in the biocontrol ability of *B. subtilis*.

## INTRODUCTION

*Bacillus subtilis* is a gram-positive bacteria belonging to the *Bacillus* genus, mainly found in soil and organic matter in decomposition (1). *B. subtilis* strains have strong colonization capacity and fast growth rate (2–4). Additionally, *B. subtilis* has been reported to produce several bioactive metabolites, such as lipopeptides and volatile organic compounds (VOCs), with strong antifungal activities (5–9). *B. subtilis* can produce a dense biofilm, which is rich in hydrolases and proteases (10,11). These interesting properties of *B. subtilis* have been effectively applied for controlling plant fungal diseases. Currently, *B. subtilis* is considered as one of the main alternatives in crop protection to avoid the use of toxic synthetic fungicides (12,13). Several biocontrol agents based on *B. subtilis*, such as Milagrum Plus (*B. subtilis* strain IAB/BS03; Idai Nature), Ethos Elite LFR (*B. subtilis* strain RTI477; FMC Corporation), Theia fungicide (*B. subtilis* strain AFS032321; AgBiome Innovations), and Minuet (*B. subtilis* strain QST713; Bayer CropScience), have been commercialized during recent years.

Black spot disease, which is caused by the fungal pathogen *Ceratocystis fimbriata*, is one of the most important diseases affecting sweet potato production and storage worldwide (14–17). The annual losses caused by black spot disease have been estimated in 30% of the global sweet potato production, leading to huge economic losses (18). Although black spot disease has been controlled during decades using synthetic fungicides, such as carbendazim, tebuconazole, and trifloxystrobin (19–21), some resistant strains have been reported (22), and the fungicides commonly used for black spot disease control have been associated to environmental pollution and major human diseases (23,24). This situation has stimulated the development of alternative strategies for black spot disease management (17). Various bacterial biocontrol agents, such as *Bacillus tequilensis* XK29 (25), *Bacillus velezensis* MMB-51 (26), *Pantoea dispersa* RO-21 (27), *Pseudomonas chlororaphis* subsp. *aureofaciens* SPS-41 (28), *Rhodococcus* sp. KB6 (29), *Streptomyces lavendulae* SPS-33 (30), and *Streptomyces setonii* WY228 (31), have been used for the control of sweet potato black rot disease. Among the reported biocontrol approaches, one of the highest efficacies was reported when using *B. subtilis* 168, which reduced black spot disease incidence and lesion length by 80% and 93%, respectively (8).

Dipicolinic acid (DPA) is one of the main components of *Bacillus* spores and plays a key role in the resistance of dormant spores to environmental stresses, such as heat, UV radiation, dryness, extreme pH, and toxic chemicals (32,33). The DPA synthase in *B. subtilis* is encoded in the *spoVF* operon, which is composed by two subunits, *spoVFA* and *spoVFB*. It has been predicted that the *spoVFA* subunit encodes the putative dehydrogenase, which is responsible for the conversion from dihydrodipicolinic acid (DHD) into DPA, while the *spoVFB* subunit encodes a flavoprotein (34). However, their actual roles in DPA biosynthesis remain unexplored. During spore germination, DPA is released from the spores through the SpoVA channels, inducing water uptake (35). DPA was identified in the fermentation broth of the widely used model strain *B. subtilis* 168 and showed strong antifungal activity, inhibiting the mycelial growth of various plant fungal pathogens (7,8,36). *B. subtilis* 168 is not able to produce lipopeptides and (37), thus, DPA plays an essential role in *B. subtilis* 168 antifungal properties. However, the transcription factor *σ^k^*, which regulates the promoter of *spoVF*, is only present in the spores, leading to low DPA contents in the vegetative cells (38). Apart from the secretion of antifungal metabolites, an appropriate attachment of the bacterial biocontrol agent onto the host surface is also an important pattern to inhibit efficiently the advancement of fungal pathogens (39). For this reason, studies to understand the key factors involved in *B. subtilis* attachment ability and biofilm formation are especially important to enhance *B. subtilis* biocontrol properties (40).

In a previous study, the promoter of *spoVF* was replaced in *B. subtilis* 168 by the promoter of *spoVG*, which is regulated by *σ^H^*, a transcription factor that is expressed in *B. subtilis* vegetative cells (8). In addition to the improved DPA production, the mutant strain also showed enhanced growth rate and biofilm formation. This report confirmed for the first time that the overexpression of the *spoVF* operon can regulate biofilm formation. However, the mechanism used by the *spoVF* operon to regulate biofilm formation is not clear. In this study, the DPA synthase genes *spoVFA* and *spoVFB* were knocked out, individually (*Δ*A and *Δ*B), or together (*Δ*A*Δ*B), in *B. subtilis* 168. The mutant strains were used to investigate the effects of the *spoVF* operon subunits on *B. subtilis* DPA synthesis, host attachment, and biofilm formation, and to clarify the regulation mechanism between the *spoVF* operon and biofilm-related genes.

## MATERIALS AND METHODS

### General information and strains

All reagents and chemicals were used as received from commercial suppliers, without further purification or modification. *B. subtilis* 168 and mutant strains were cultured in lysogeny broth (LB; 5 g yeast extract, 10 g peptone, and 10 g sodium chloride in 1 L of distilled water, pH 7.0-7.2) at 37 °C (7). LB semisolid medium was prepared after adding 1.5 g/L agar. The construction of the mutant strain with enhanced *spoVF* expression by replacing the original promoter by *spoVG* promoter was performed as previously reported (8).

Carbendazim-resistant *C. fimbriata* isolate NJC (GenBank accession numbers: MT560374.1 and PQ232785.1), which was previously isolated from sweet potatoes with black spot disease symptoms (14,22), was used in this study. This fungal strain was cultured on potato-glucose-agar (PDA) medium, which was prepared by boiling 200 g potatoes in 1 L ddH_2_O for 30 min, and then adding 20 g glucose (pH = 5.6) (41). Sweet potatoes (*Ipomoea batatas* L. Lam) were purchased from a local supermarket. A ZQZY-78BV oscillating incubator (Shanghai Zhichu, China) was used for bacterial fermentation.

### Construction of the knockout strains

The construction of the knockout strains was carried out using the homologous recombination method previously reported (8). The target genes were *spoVFA* (GenBank number: 939652) and *spoVFB* (GenBank number: 939660). The homologous recombination for *Δ*A construction was carried out by overlapping the extension fragment that consisted of the upstream of *spoVFA* gene (498 bp), chloramphenicol expression cassette, and downstream of *spoVFA* gene (502 bp). Similarly, for *Δ*B construction, the overlapping extension fragment consisted of the upstream of *spoVFB* gene (499 bp), chloramphenicol expression cassette, and downstream of *spoVFB* gene (501 bp). The overlapping extension fragment for *Δ*A*Δ*B construction consisted of the upstream of *spoVFA* gene (498 bp), chloramphenicol expression cassette, and downstream of *spoVFB* gene (501 bp). LB containing chloramphenicol (5 μg/mL) was used to select the positive clones. Primers used for the construction of the mutant strains are shown in Supporting Information Table S1. The amplified fragments are shown in Supporting Information Table S2. In this study, WT stands for wildtype *B. subtilis* strain 168, MT stands for the mutant with enhanced *spoVF* operon expression obtained after promoter replacement (8), *Δ*A stands for the *spoVFA* knockout strain, *Δ*B stands for the *spoVFB* knockout strain, and *Δ*A*Δ*B stands for the *spoVFA* and *spoVFB* double knockout strain.

### Gram staining of *B. subtilis* strains

The five strains used in this study (WT, MT, *Δ*A, *Δ*B, and *Δ*A*Δ*B) were mixed with sterile ddH_2_O on 25 × 75 mm clean slides and spread to form a thin and uniform round membrane of 10 ∼ 15 mm diameter. After drying at room temperature, 20 µL crystal violet (the solution was prepared by dissolving 2 g crystal violet in 20 mL 95 % ethanol and 80 mL 1% ammonium oxalate aqueous solution) (crystal violet was purchased from Shanghai Yuanhang, China) was used to stain the cells for 1 min. After washing with H_2_O until no stain was observed in the discarded solution, 20 µL Lugol’s iodine solution (0.033 g/L, Macklin, China) was used to stain the cells for 1 min. Then, the cells were washed with H_2_O until no stain was observed in the discarded solution. Ethanol (95%) was used to decolor the dye for 20 ∼ 30 s, and then the cells were washed with H_2_O immediately. Finally, 20 µL saffron (2.5 g saffron in 100 mL 95% ethanol) (saffron was purchased from Shanghai Tianhuang, China) was used to stain the cells for 3 ∼ 5 min. After washing with H_2_O until no stain was observed in the discarded solution and drying at room temperature, cell morphology was were observed using a DM2500 microscope (Leica, Japan).

### Analysis of cell growth and DPA concentration

The five strains (WT, MT, *Δ*A, *Δ*B, and *Δ*A*Δ*B) were pre-cultured in LB medium (5 mL) at 37 °C and 200 rpm overnight. Seed cultures were grown at 37 °C and 200 rpm in 500 mL Erlenmeyer flasks containing 50 mL LB medium until OD_600_ = 2.0. Aerobic fermentations were carried out in 500 mL-baffled Erlenmeyer flasks containing 50 mL LB medium with 20 g/L glucose at 37 °C and 200 rpm using an initial OD_600_ of 0.1. Cell growth was monitored at 600 nm (OD_600_) using a BioMate 3 spectrophotometer (722S, Lengguang Technology, China). The culture was diluted 20 times with ddH_2_O to ensure that the measured OD_600_ value was in the range from 0.2 to 0.8. Glucose concentration in the culture medium was measured using the 3,5-dinitrosalicylic acid (DNS) method (42). Three independent assays were performed.

DPA concentration was detected after 8, 12, 16, and 20 h fermentation. To achieve this goal, 5 mL of *B. subtilis* fermentation culture was centrifuged at 10,000 rpm and 4 °C for 6 min. Then, the filtered supernatant was examined by high-performance liquid chromatography (HPLC; Agilent 1200 series, Hewlett–Packard, USA) with an UV-visible light absorbance detector (at 254 nm) and a C-18 column (250 × 3.0 mm, Phenomenex) at 35 ℃. A mixture of methanol/acetic acid/ddH_2_O (12.5:1.5:86.0 *v/v/v*) was used as the mobile phase at a constant flow rate of 0.4 mL/min for 15 min (41). The retention time of DPA using the mentioned conditions was 7.2 min. Commercial DPA (Macklin, China) was used as the standard. Three independent assays were performed.

### Biofilm formation analysis

The five strains (WT, MT, *Δ*A, *Δ*B, and *Δ*A*Δ*B) were inoculated in 10 mL LB medium and pre-cultured at 37 °C and 200 rpm to OD_600_ = 1.2. The cells were collected by centrifugation at 5,000 rpm and 4 °C for 8 min, and then resuspended in an equal volume of sterile ddH_2_O. A 30 µL aliquot of cell suspension was then inoculated into 3 mL LB medium, in a glass test tube, and incubated at 37 °C for 3 days without shaking. After removing the aqueous solution, 6 mL 10% crystal violet solution (dissolved in 95 % ethanol) was added to the glass tube, and the biofilm was treated for 1 h. Then, the test tube was washed three times with 10 mL ddH_2_O and air dried for 1 h. Finally, 3 mL 40% methanol and 10% glacial acetic acid aqueous solution was added to the test tube to dissolve the crystal violet stain. The absorbance of the dissolved crystal violet was determined using a spectrophotometer (722S, Lengguang Technology, China) at 575 nm (43).

### Evaluation of the colonizing ability of *B. subtilis* strains on sweet potato slices

The five strains (WT, MT, *Δ*A, *Δ*B, and *Δ*A*Δ*B) were cultured in 5 mL LB medium at 37 °C and 200 rpm overnight. Then, 300 µL bacterial suspension was inoculated into 50 mL LB medium and cultured at 37 °C and 200 rpm for 6 h. After centrifugation at 8,000 rpm and 4 °C for 6 min, the cells were collected and washed with 5 mL sterile ddH_2_O. Finally, the cells were resuspended in 20 mL sterile ddH_2_O to OD_600_ = 1.2. The colonization ability of the strains was monitored by scanning electron microscopy (SEM) and population dynamics using previously reported procedures (8). Sweet potato slices, 1 cm in width, were cut and washed with 10 mL 5% NaClO for 15 min. After discarding the NaClO solution, the sweet potato slices were washed with 10 mL sterile ddH_2_O three times. Then, 10 mL cell suspensions (OD_600_ = 1.2; 4.4 × 10^5^ CFU/mL) were sprayed onto the sweet potato slices. Six sweet potato slices were used for each strain (10 mL cell suspension was sprayed on six sweet potato slices). After drying for 30 min, a 1 cm × 1 cm piece was cut off from each slide and carefully washed with 1 mL ddH_2_O. SEM analysis of the sweet potato surface was carried out using a Gemini 300 instrument (Zeiss, USA). The preparation of the SEM samples was carried out as previously reported (8). The number of cells on the sweet potato samples was correlated with the attachment capacity of the strains.

On the other hand, the washing solution was collected and diluted 4,000 times. Then, 100 µL of the resulting solution was coated on a 9-cm-diameter LB plate, which were then cultured at 37 °C overnight. The concentration of bacteria was calculated in terms of colony forming units per plate (CFU per plate) in the washing solution (8). The higher concentration of bacteria in the washing solutions was inversely correlated with the attachment ability of the bacterial strains.

### Detection of mRNA levels of key genes

The five strains (WT, MT, *Δ*A, *Δ*B, and *Δ*A*Δ*B) were inoculated in 5 mL LB medium and cultured at 37 °C and 200 rpm overnight. The cultures were centrifuged at 4 ℃ and 10,000 rpm for 6 min. Then, the supernatant was discarded, and the cell pellets were washed twice with 5 mL sterile ddH_2_O. Total RNA was extracted with TRIzol reagent (Thermo Fisher Scientific, China). The remaining DNA was removed with the Transcript All-in-One-first-strand DNA Synthesis SuperMix for qPCR (one Step gDNA Removal) Kit (Vazyme Biotech, China). cDNA was synthesized by reverse transcription, and qRT-PCR was carried out with the SYBR Green I Real-Time PCR Kit (Solarbio, China). A 7500 Realtime PCR system (Applied Biosystems, ABI, USA) was used for the PCR analysis. *16S rRNA* was used as the internal reference gene (8), and the relative expression was calculated by the 2^-ΔΔCt^ method. The tested genes, which are related to DPA biosynthesis and tricarboxylic acid (TCA) cycle, and the primers used in the qRT-PCR analysis are shown in Supporting Information Table S3.

### Detection of NADP(H) content

The NADP^+^/NADPH Test Kit (Beyotime Biotechnology, China) was used to detect the contents of NADP^+^ and NADPH (44). The five strains (WT, MT, *Δ*A, *Δ*B, and *Δ*A*Δ*B) were cultured in LB medium (5 mL) at 37 ℃ and 200 rpm for 12 h. One milliliter of cell suspensions (containing 1 × 10^6^ CFU/mL) were collected and centrifuged at 6,000 rpm for 5 min. After discarding the supernatant, 200 μL ice pre-cooled NADP^+^/NADPH extraction solution was added and gently mixed to promote cell lysis. Subsequently, the mixture was centrifuged at 12,000 rpm and 4 °C for 10 min, and the supernatant was collected. The manufacturer’s instructions were followed to calculate the amounts of NADP^+^ and NADPH in the cells. The principle followed by this kit relies on the glucose-6-phosphate dehydrogenase-catalyzed conversion from glucose-6-phosphate into D-glucono-1,6-lactone in the presence of NADP^+^, and the transformation from 2-(2-methoxy-4-nitrophenyl)-3-(4-nitrophenyl)-5-(2,4-disulfophenyl)-2H-tetrazolium into formazan in the presence of 1-methoxy-5-methylphenazinium methyl sulfate and NADPH. Thus, different detectable products are formed depending on the concentrations of NADP^+^ and NADPH.

### Evaluation of *B. subtilis* stress tolerance

The five strains (WT, MT, *Δ*A, *Δ*B, and *Δ*A*Δ*B) were grown in LB medium (5 mL) at 37 ℃ and 200 rpm to OD_600_= 1.2. The tolerance of the strains to different temperatures, NaCl concentrations, pH values, and UV light was examined. To study the tolerance of the strains to high temperature, the cell suspensions were diluted 10^6^ times with sterile ddH_2_O for the further analysis. Then, 100 µL aliquots were coated on 9-cm-diameter Petri dishes containing LB agar medium, which were subsequently incubated at 30, 37, and 42 ℃ for 24 h (46). The bacterial survival was measured according to the CFU per plate.

To study the tolerance of the strains to osmotic pressure, 1 mL cell suspensions (4.4 × 10^5^ CFU/mL) were centrifuged at 8,000 rpm and 4 ℃ for 4 min. After discarding the supernatant, the cells were spread on 9-cm-diameter Petri dishes, containing LB semisolid medium with 0, 9, 18, 36, and 54 g/L NaCl. The cultures were incubated at 37 ℃ and 200 rpm for 24 h. The bacterial survival was measured according to the CFU per plate.

To study the tolerance of the strains to acidic and alkaline pH values, 1 mL cell suspensions (4.4 × 10^5^ CFU/mL) were centrifuged at 8,000 rpm and 4 ℃ for 4 min. After discarding the supernatant, the cells were resuspended in 1 mL LB medium with pH values ranging from 4 to 9. The resulting suspensions were cultured at 37 ℃ and 200 rpm for 24 h. Then, 100 µL cell culture was spread on 9-cm-diameter Petri dishes containing LB semisolid medium, and the resulting cultures were incubated at 37 ℃ for 24 h. The survival was monitored according to the CFU on each plate. Each treatment was repeated three times.

To study the tolerance of the strains to UV light, 1 mL cell suspensions were centrifuged at 8,000 rpm and 4 ℃ for 4 min. After discarding the supernatant, the cells were resuspended in 1 mL sterile ddH_2_O, and 100 µL of the resulting cell suspension was coated on a 9-cm-diameter Petri dish containing LB semisolid medium. Then, UVA (340 nm) and UVB (313 nm) irradiations were applied on the Petri dishes for 5 and 20 min using a UVA lamp (T8 20W, UVA-340 nm, Yixian, China) and a UVB lamp (G20T12E, UVB-313 nm, Yixian, China). The CFU per plate was counted after culturing at 37 ℃ for 24 h. Each treatment was repeated 3 times.

### In vitro antifungal activity of B. subtilis against C. fimbriata

The five strains (WT, MT, *Δ*A, *Δ*B, and *Δ*A*Δ*B) were pre-incubated in LB medium (5 mL) at 37 °C and 200 rpm to OD_600_ = 1.2. Cells were harvested via centrifugation at 5,000 rpm and 4 °C for 8 min, and resuspended in 1 mL ddH_2_O to OD_600_ = 1.2. A plug of *C. fimbriata* was located on the center of a 9-cm-diameter PDA plate, and 6 µL bacterial strain solution was spread into a straight line on the left side of the plug. The distance between the plug and the straight line containing the bacterial strains was 2 cm. The PDA plates containing *C. fimbriata* and the bacterial strains were incubated at 28 °C for 72 h. The antifungal activity was measured according to the inhibition of the mycelial growth. The control experiment was carried out by culturing *C. fimbriata* in the absence of bacterial strains. To check the effects of the bacterial strains on the viability of *C. fimbriata* cells, Neutral red (0.1 mg/mL) (Beyotime Biotechnology, China) and Evans blue (0.5 mg/mL) (Aladdin, USA) were used to stain the fungal cells following the procedure previously reported by Jiang et al. (27). Briefly, 10 µL of the dye solution was dropped on a 25 mm × 75 mm glass slide. Then, the fungal cells, which were collected from different locations of the plate, near and far away from the bacterial strains, were dipped in the dye solutions, incubated at room temperature for 5 min, and washed 3 times with sterile ddH_2_O (1 mL). Each treatment was repeated 3 times. The stained cells were observed using an DM2500 microscope (Leica, Japan).

### Assessment of competition for nutrients

The effect of nutrient depletion by the five strains (WT, MT, *Δ*A, *Δ*B, and *Δ*A*Δ*B) on *C. fimbriata* spore germination was tested following the conditions previously reported by Di Francesco et al. (45). The reported method used two polystyrene cylinders divided by a hydrophilic polytetrafluoroethylene (PTFE) membrane, with a pore size of 0.45 μm, which allowed the exchange of nutrients but not cells. Sweet potato juice was obtained by boiling 100 g of sweet potato pulp for 20 min in 500 mL ddH_2_O. The cylinders were filled with 0%, 0.5%, and 5% sterilized sweet potato juice in ddH_2_O (25 mL). In one cylinder, 250 μL *C. fimbriata* spore solution (1 × 10^5^ spores/mL) was added, while 250 μL *B. subtilis* cell solution (1 × 10^3^ CFU/mL) was added in the other cylinder. The control experiment was carried out by adding a *C. fimbriata* spore suspension (1 × 10^5^ spores/mL) to one cylinder, in the absence of biocontrol strains in the second cylinder (41). The cylinders were placed at 28 ℃ on a rotary shaker at 120 rpm. After 24 h incubation, suspensions were collected and transferred to a glass slide for microscope observations. An Olympus BX43 Upright Microscope (Germany) was used to calculate the ratio of germinated spores. A spore was considered germinated when the germination tube was at least 50% in length compared to the total length of the spore.

### *In vivo* efficacy of *B. subtilis* for the control of black rot disease

*C. fimbriata* was cultured on PDA medium for 5 days, and the mycelium was divided into 5-mm-diameter plugs. Three plugs were cultured in 10 mL PDA broth (without agar) at 28 °C and 200 rpm for 2 days. After centrifuging at 4 ℃ and 5,000 rpm for 6 min, the supernatant was discarded, and the collected spores were resuspended in ddH_2_O to 1 × 10^5^ spores/mL concentration. The five strains (WT, MT, *Δ*A, *Δ*B, and *Δ*A*Δ*B) were grown in 5 mL LB medium at 37 °C and 200 rpm to OD_600_ = 5.0 (1.4 × 10^8^ CFU/mL). After centrifuging at 4 ℃ and 5,000 rpm for 6 min, bacterial cells were washed twice with 1 mL ddH_2_O, and finally re-suspended in a suitable amount of sterilized ddH_2_O to 1 × 10^7^ and 1 × 10^8^ CFU/mL. Sweet potatoes were cut into slices, of approximately 6 cm diameter and 4 cm width. Two 1-cm-diameter holes, approximately 1 cm in depth, were dug in the outer edge of the cross section of each slice (47). Twenty sweet potato slices were used for each treatment condition. To study the antifungal activity of the bacterial strains, 20 mL bacterial suspension (containing 1 × 10^7^ or 1 × 10^8^ CFU/mL) was sprayed on the twenty sweet potato slices. For the negative control experiment, the same volume of sterilized ddH_2_O was sprayed. After drying, 15 μL *C. fimbriata* aqueous suspension (1 × 10^5^ spores/mL) was injected in each hole, and the sweet potato slices were incubated at 28 °C and 80% relative humidity. After 72 h, the pathogen advancement was calculated according to the lesion length caused by *C. fimbriata*. The lesions showed brown/black color and could be easily observed with the naked eye (8). Four independent assays, with twenty sweet potatoes for each treatment condition in each one, were performed.

## Data analysis

SPSS (statistical Package, Version 20.0) software was used for statistical analysis. Variables were tested for significance using the Student’s t-test at the *p* < 0.05 (*), *p* < 0.01 (**), *p* < 0.001 (***), and *p* < 0.0001 (****) levels (ns = no significance).

## RESULTS

### Roles of *spoVF* subunits in *B. subtilis* morphology and growth

Microscope observations indicated the presence of purple color cells, which is in agreement with the Gram-positive nature of *B. subtilis* (Fig. 1). Both WT and MT strains showed rod-shaped and were 2.72 ± 0.48 µm in length. Surprisingly, some *Δ*A, *Δ*B, and *Δ*A*Δ*B cells were spherical. This demonstrates that the *spoVF* operon plays a key role in *B. subtilis* morphology. It is worth mentioning that the proportion of spherical cells was higher in *Δ*A (100% spherical cells) than in *Δ*B (7.1% spherical cells). The proportion of *Δ*A*Δ*B spherical cells was 100%.

**FIG 1.**
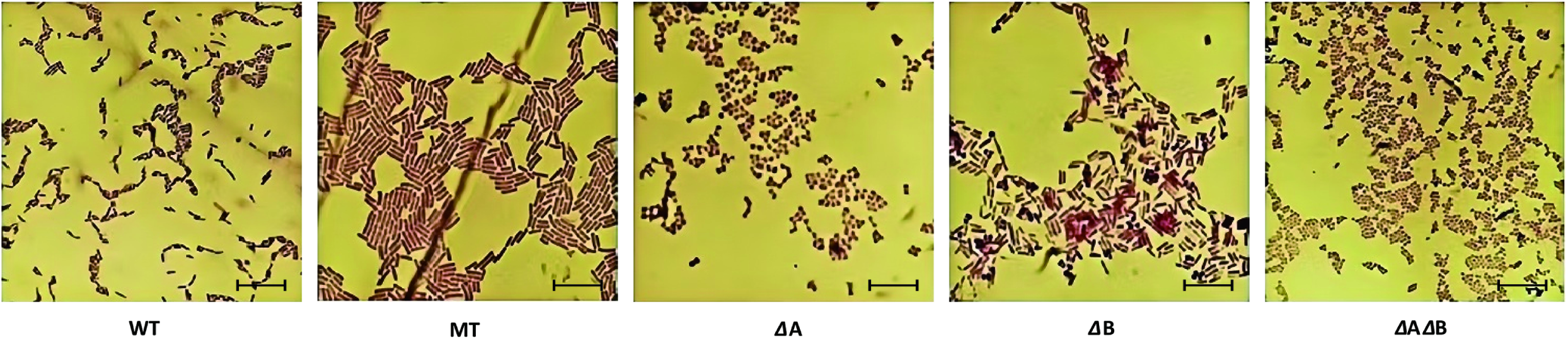
Gram staining results of different *Bacillus subtilis*. *Abbreviations:* WT, wildtype *B. subtilis* strain 168; MT, mutant obtained after promoter replacement (8); *Δ*A, *spoVFA* knockout strain; *Δ*B, *spoVFB* knockout strain; *Δ*A*Δ*B, *spoVFA* and *spoVFB* double knockout strain. Scale bar = 10 µm.

Consistently with a previous study (8), the growth of MT strain was faster than the growth of WT strain (Fig. 2A). In agreement, the glucose consumption rate of MT strain was higher than that of WT (Fig. 2B). Cell growth and glucose consumption of *Δ*A and *Δ*B strains were similar between them, and higher compared to those of *Δ*A*Δ*B strain. Cell growth and glucose consumption of *Δ*A and *Δ*B strains were lower compared to those of the WT strain. The OD_600_ value in the fermentation broth of MT, WT, *Δ*A, *Δ*B, and *Δ*A*Δ*B after 16 h was 13.8 ± 0.6, 11.0 ± 0.5, 8.9 ± 0.5, 8.6 ± 0.3, and 8.3 ± 0.3, respectively, whereas glucose consumption by MT, WT, *Δ*A, *Δ*B, and *Δ*A*Δ*B strains after 24 h was 83.5%, 82.5%, 76.5%, 78.0%, and 73.5%, respectively. This indicates that both *spoVF* operon subunits are involved in *B. subtilis* growth and glucose consumption.

**FIG 2.**
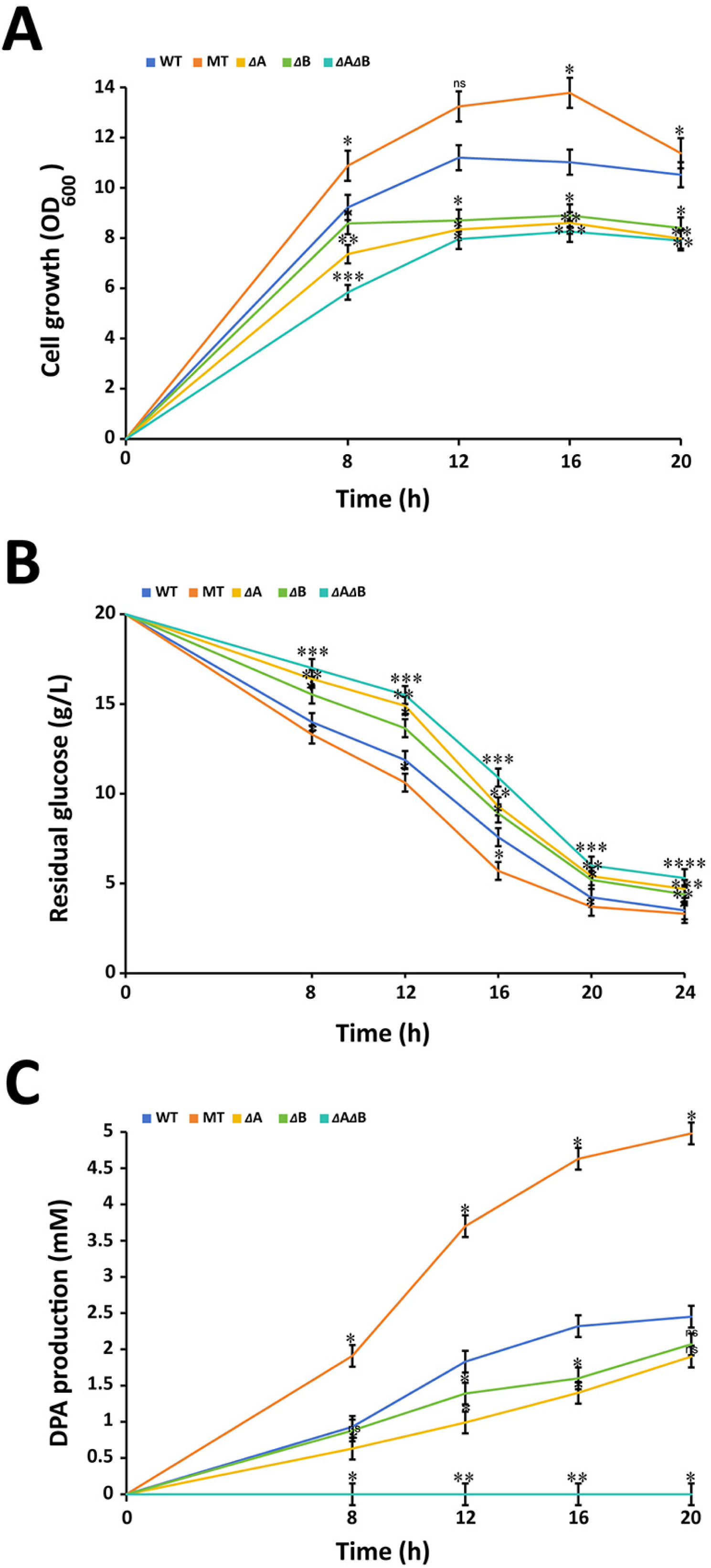
Effects of *spoVF* operon subunits on *Bacillus subtilis* growth, glucose consumption, and DPA production. **(A)** Cell growth (measured according to the OD_600_ value). Lysogeny broth (LB) medium with 20 g/L glucose was used for the fermentation. **(B)** Glucose consumption. Glucose concentration was measured using the 3,5-dinitrosalicylic acid (DNS) method (42). **(C)** Dipicolinic acid (DPA) production after 8, 12, 16, and 20 h of fermentation. High-performance liquid chromatography (HPLC; Agilent 1200 series, Hewlett–Packard, USA) with an UV-visible light absorbance detector (at 254 nm) and a C-18 column (250 × 3.0 mm, Phenomenex) was used for the detection. A mixture of methanol/acetic acid/ddH_2_O (12.5:1.5:86.0 *v/v/v*) was used as the mobile phase at a constant flow rate of 0.4 mL/min for 15 min (41). *Abbreviations:* WT, wildtype *B. subtilis* strain 168; MT, mutant obtained after promoter replacement (8); *Δ*A, *spoVFA* knockout strain; *Δ*B, *spoVFB* knockout strain; *Δ*A*Δ*B, *spoVFA* and *spoVFB* double knockout strain. Means were submitted to Student’s t test, with significance values at the *p* < 0.05 (*), *p* < 0.01 (**), *p* < 0.001 (***), and *p* < 0.0001 (****) levels (ns = no significance). Means at different times were compared with those of WT.

DPA yield was higher when using the MT strain than the WT strain (Fig. 2C). This can be explained considering that MT contains a new promoter that can express the operon in the vegetative cells. Although WT, *Δ*A, and *Δ*B strains did not have the replaced promoter in the *spoVF* operon, low concentrations of DPA were still detected in their fermentation medium. Among the three strains, DPA production with WT strain was the highest (2.45 ± 0.2 m/L DPA after 24 h), followed by the *Δ*B strain (2.07 ± 0.2 m/L DPA after 24 h), and then the *Δ*A strain (1.9 ± 0.2 m/L DPA after 24 h). DPA could not be detected in fermentation medium of the *Δ*A*Δ*B strain. This suggests that the two subunits of the *spoVF* operon are involved in DPA synthesis. However, the knockout of both subunits together is necessary to completely eliminate the ability to produce DPA. These results also suggest that DPA can be synthesized by both subunits independently. However, subunit *spoVFA* seems to be more important regarding DPA biosynthesis than the subunit *spoVFB*.

### Roles of *spoVF* subunits in *B. subtilis* biofilm formation and colonization ability on sweet potato

The OD_575_ values of the crystal violet-stained biofilm produced by WT, MT, *Δ*A, *Δ*B, and *Δ*A*Δ*B strains were 9.56 ± 0.34, 19.71 ± 0.92, 3.76 ± 0.06, 7.47 ± 0.01, and 2.58 ± 0.16, respectively (Fig. 3). Thus, the MT strain exhibited the highest biofilm formation ability, followed by WT, *Δ*B, *Δ*A, and *Δ*A*Δ*B. This indicates that biofilm formation ability is positively correlated with the expression level of *spoVF* operon. The trend in biofilm production was proportional to the trends in growth, glucose consumption, and DPA production ability, suggesting that the four factors are related between them.

**FIG 3.**
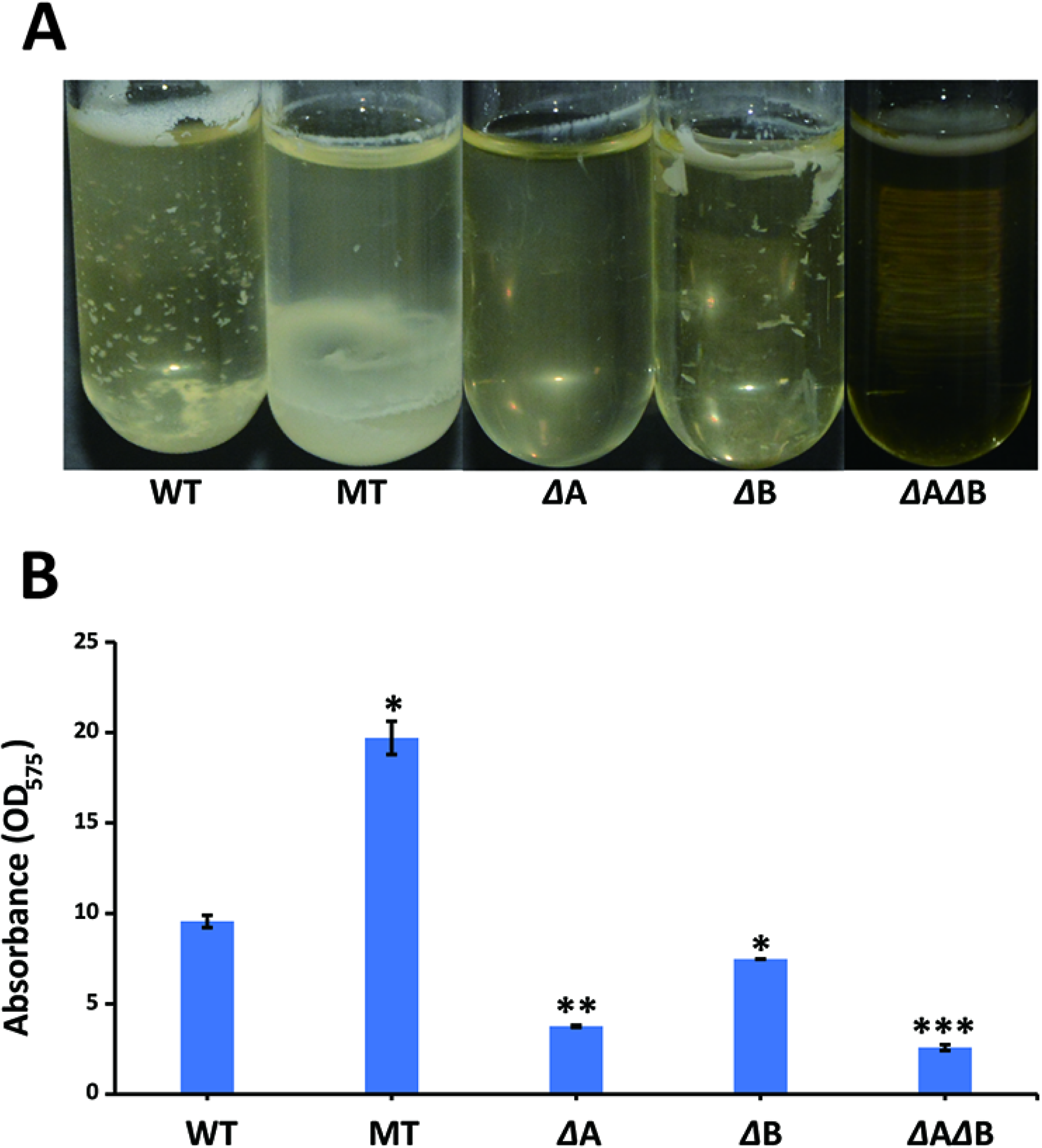
Effects of *spoVF* operon subunits on *Bacillus subtilis* biofilm formation. **(A)** Images of the biofilm formation produced by the bacterial strains. **(B)** Measurement of biofilm formation by the bacterial strains. The biofilm formation was measured according to the OD_575_ value after staining with crystal violet (43). *Abbreviations:* WT, wildtype *B. subtilis* strain 168; MT, mutant obtained after promoter replacement;^8^ *Δ*A, *spoVFA* knockout strain; *Δ* B, *spoVFB* knockout strain; *Δ*A*Δ*B, *spoVFA* and *spoVFB* double knockout strain. Means at different times were compared with those of WT.

SEM images indicated that some *Δ*B, *Δ*A, and *Δ*A*Δ*B cells showed spherical shape (Fig. 4A), which is consistent with the microscope observations after Gram staining. The MT strain exhibited the highest colonization ability on sweet potato surface, followed by WT, *Δ*B, *Δ*A, and *Δ*A*Δ*B. These results are consistent with the results related to biofilm formation. This can be explained considering that biofilm formation is a key factor involved in the colonization capacity of *B. subtilis* (48). The number of CFU per plate in the washing solutions of WT, MT, *Δ*A, *Δ*B, and *Δ*A*Δ*B strains was 29.6 ± 4.3, 7.8 ± 2.7, 49.2 ± 7.8, 38.4 ± 5.1, and 74.6 ± 10.6, respectively (Fig. 4B).

**FIG 4.**
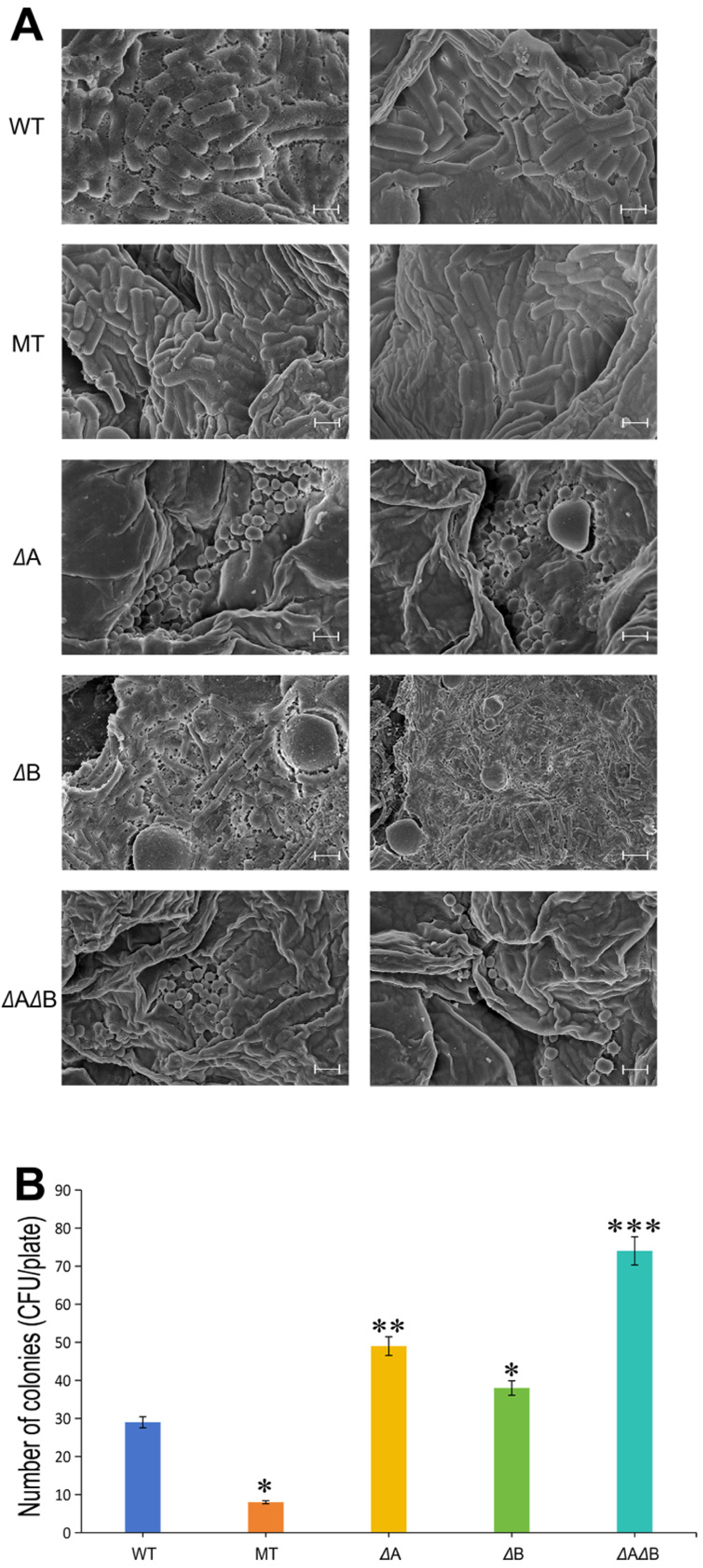
Effects of *spoVF* operon subunits on *Bacillus subtilis* colonization ability on sweet potato. **(A)** Scanning microscopy (SEM) images showing the colonization pattern of the bacterial strains on sweet potato slices. Scale bar = 2 µm. **(B)** Number of colonies detected in the sweet potato washing solution. The sweet potato slices were washed with ddH_2_O to understand the attachment ability of the bacterial strains on the sweet potato surface. *Abbreviations:* WT, wildtype *B. subtilis* strain 168; MT, mutant obtained after promoter replacement (8); *Δ*A, *spoVFA* knockout strain; *Δ*B, *spoVFB* knockout strain; *Δ*A*Δ*B, *spoVFA* and *spoVFB* double knockout strain.

These results indicated that the MT strain could attach better to the sweet potato surface in comparison with the WT strain, whereas the *Δ*B, *Δ*A, and *Δ*A*Δ*B strains showed poor colonization capacity and could be easily removed from the sweet potato surface with water.

### Roles of *spoVF* subunits in *B. subtilis* TCA cycle gene expression and NADPH/NADP^+^ biosynthesis

As expected, no expression for *spoVFA* and *spoVFB* genes was detected in *Δ*A and *Δ*B strains, respectively (Table 1). Consistently, no expression for *spoVFA* and *spoVFB* genes was detected in the *Δ*A*Δ*B strain. The expression of *asd*, *dapG*, and *dapA* genes, which are involved in the first steps of DPA biosynthesis, was downregulated in *Δ*A, *Δ*B, and *Δ*A*Δ*B strains in comparison with the WT strain. The expression levels of these genes in *Δ*A strain were significantly lower than in *Δ*B strain. These results indicated that, among the *spoVF* operon subunits, *spoVFA* plays a more important role than *spoVFB* in the activation of the DPA biosynthetic pathway. This is consistent with the different DPA production by *Δ*A and *Δ*B strains (Section 3.1.). In contrast, *spoVFA*, *spoVFB*, *asd*, *dapG*, and *dapA* genes were upregulated in MT.

**TABLE 1.** qRT-PCR analysis of key genes in the bacterial strains.^a^.

| Gene categories | Gene name | Gene expression fold change compared to WT<sup>b</sup> |  |  |  |
| --- | --- | --- | --- | --- | --- |
| | | MT | $\Delta A$ | $\Delta B$ | $\Delta A\Delta B$ |
| DPA synthesis pathway | *spoVFA* | 8.60 ± 0.23 | -<sup>c</sup> | 0.90 ± 0.18 | -<sup>c</sup> |
|  | *spoVFB* | 9.10 ± 0.35 | 1.10 ± 0.20 | -<sup>c</sup> | -<sup>c</sup> |
|  | *asd* | 8.50 ± 0.21 | 0.02 ± 0.01 | 0.35 ± 0.03 | 0.02 ± 0.01 |
|  | *dapG* | 5.30 ± 0.30 | 0.18 ± 0.02 | 0.22 ± 0.02 | 0.47 ± 0.11 |
|  | *dapA* | 7.90 ± 0.28 | 0.02 ± 0.01 | 0.80 ± 0.01 | 0.09 ± 0.03 |
|  | *pdhA* | 2.70 ± 0.13 | 0.31 ± 0.02 | 0.36 ± 0.08 | 0.35 ± 0.06 |
| TCA cycle | *citC* | 4.72 ± 0.34 | 0.19 ± 0.02 | 0.49 ± 0.06 | 0.32 ± 0.07 |
|  | *gltA* | 2.66 ± 0.24 | 0.03 ± 0.01 | 0.017 ± 0.002 | 0.73 ± 0.12 |
| Transcription factor | *spo0A* | 7.59 ± 0.82 | 0.20 ± 0.01 | 0.25 ± 0.01 | 0.15 ± 0.03 |
| Biofilm formation | *kinC* | 1.80 ± 0.16 | 0.06 ± 0.01 | 0.13 ± 0.01 | 0.68 ± 0.05 |
|  | *sinR* | 0.10 ± 0.02 | 31.96 ± 3.71 | 25.50 ± 1.72 | 16.77 ± 0.79 |
|  | *abrB* | 0.21 ± 0.02 | 4.34 ± 0.49 | 2.02 ± 0.10 | 1.84 ± 0.08 |
|  | *tapA* | 2.16 ± 0.11 | 0.16 ± 0.04 | 0.22 ± 0.07 | 0.53 ± 0.09 |
| Cell morphology | *merB1* | 4.13 ± 0.36 | -<sup>c</sup> | -<sup>c</sup> | -<sup>c</sup> |
|  | *merB2* | 4.14 ± 0.17 | -<sup>c</sup> | -<sup>c</sup> | -<sup>c</sup> |

<sup>a</sup>The primers used for the qRT-PCR assay are indicated in Supporting Information Table S3.<sup>b</sup>The expression of the genes in WT was adjusted to 1.<sup>c</sup>Not detected.
Abbreviations: WT, wildtype *B. subtilis* strain 168; MT, mutant obtained after promoter replacement (8); $\Delta A$ , *spoVFA* knockout strain; $\Delta B$ , *spoVFB* knockout strain; $\Delta A\Delta B$ , *spoVFA* and *spoVFB* double knockout strain.

Pyruvate dehydrogenase, which is encoded by *pdhA*, catalyzes the reaction from pyruvate to acetyl-CoA, which is the substrate of the first reaction of the TCA cycle. The expression level of *pdhA* was 2.7-fold higher in the MT strain in comparison with that in the WT strain (Table 1). Isocitrate dehydrogenase, which is encoded by *citC*, catalyzes the reaction from isocitrate into α-ketoglutarate, with the simultaneous reduction of NADP^+^ into NADPH. The expression level of *citC* was 4.7 times higher in MT than in WT. Glutamate synthase (GOGAT), which is encoded by *gltA*, catalyzes the transformation from α-ketoglutaric acid to glutamic acid. The expression level of *gltA* was 2.7 times higher in MT than in WT. Spo0A is a transcription factor of *B. subtilis*, and KinC can catalyze Spo0A phosphorylation. The expression of levels of *gltA* and *kinC* increased by 7.6-and 1.8-fold, respectively, in MT strain compared to those in WT strain. The extracellular polysaccharide TasA amyloid fiber, which is synthesized by the *epsA* and *tapA* operons, is the main component of *B. subtilis* biofilm. The transcription factors AbrB and SinR can inhibit *epsA* and *tapA* expression. Compare with WT strain, the expression level of *tapA* was 2.2 times higher in MT, and *sinR* and *abrB* expression levels were 10.0 and 4.7 times lower, respectively. The upregulation and downregulation trends observed in the expression levels of *pdhA*, *citC*, *gltA*, *kinC*, *tapA*, *sinR*, and *abrB* in *Δ*A, *Δ*B, and *Δ*A*Δ*B strains were exactly the opposite than those observed in MT.

The expression of *merB1* and *merB2*, which encode cell-shape determining proteins, was significantly downregulated in *Δ*A, *Δ*B, and *Δ*A*Δ*B compared to WT (Table 1). These results are consistent with the changes in cell morphology in those strains (Section 3.1.) and confirm that the *spoVF* operon plays a key role in *B. subtilis* morphology.

As it can be seen in Fig. 5, MT contained the highest NADPH and NADP^+^ levels during fermentation. When increasing the fermentation time, NADPH concentration increased in MT, whereas NADPH concentration decreased in the other four strains. The concentration of NADP^+^ decreased over time in the fermentation medium of the five strains.

**FIG 5.**
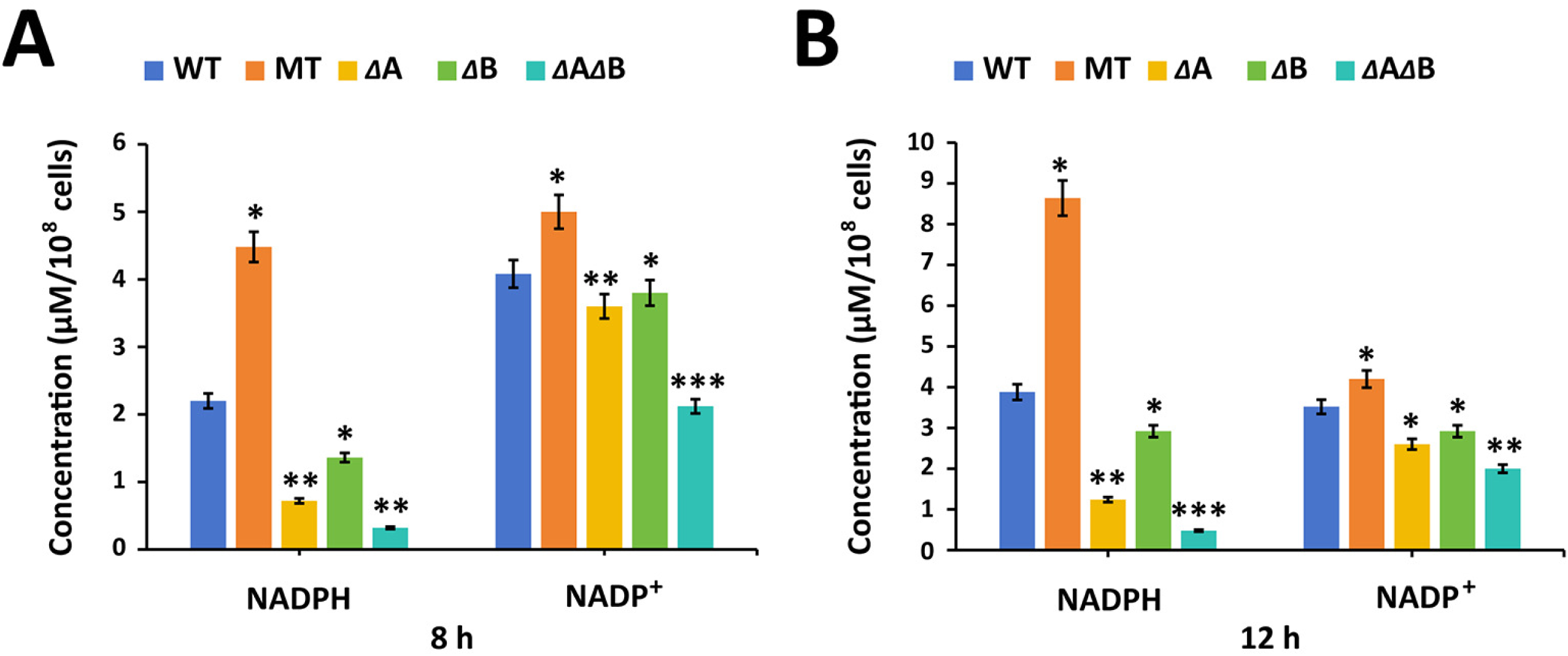
Effects of *spoVF* operon subunits on the contents of intracellular NADPH and NADP^+^. **(A)** Contents of NADPH and NADP^+^after 8 h fermentation. **(B)** Contents of NADPH and NADP^+^after 12 h fermentation. *Abbreviations:* WT, wildtype *B. subtilis* strain 168; MT, mutant obtained after promoter replacement (8); *Δ*A, *spoVFA* knockout strain; *Δ*B, *spoVFB* knockout strain; *Δ*A*Δ*B, *spoVFA* and *spoVFB* double knockout strain. Means were submitted to Student’s t test, with significance values at the *p* < 0.05 (*), *p* < 0.01 (**), *p* < 0.001 (***), and *p* < 0.0001 (****) levels (ns = no significance). Means at different times were compared with those of WT.

### Roles of *spoVF* subunits in *B. subtilis* tolerance to stress conditions

As shown in Fig. 6A, WT, MT, *Δ*B, *Δ*A, and *Δ*A*Δ*B showed the highest CFU values at 37 ℃, whereas the lowest CFU values were observed at 42 ℃. This suggests that 42 ℃ challenges bacterial survival. The CFU values of MT were the highest at all different temperatures, followed by WT, *Δ*B, *Δ*A, and *Δ*A*Δ*B. Both *Δ*A and *Δ*A*Δ*B did not grow at 42 ℃, indicating that the *spoVFA* subunit is a key factor involved in *B. subtilis* tolerance to heat stress.

**FIG 6.**
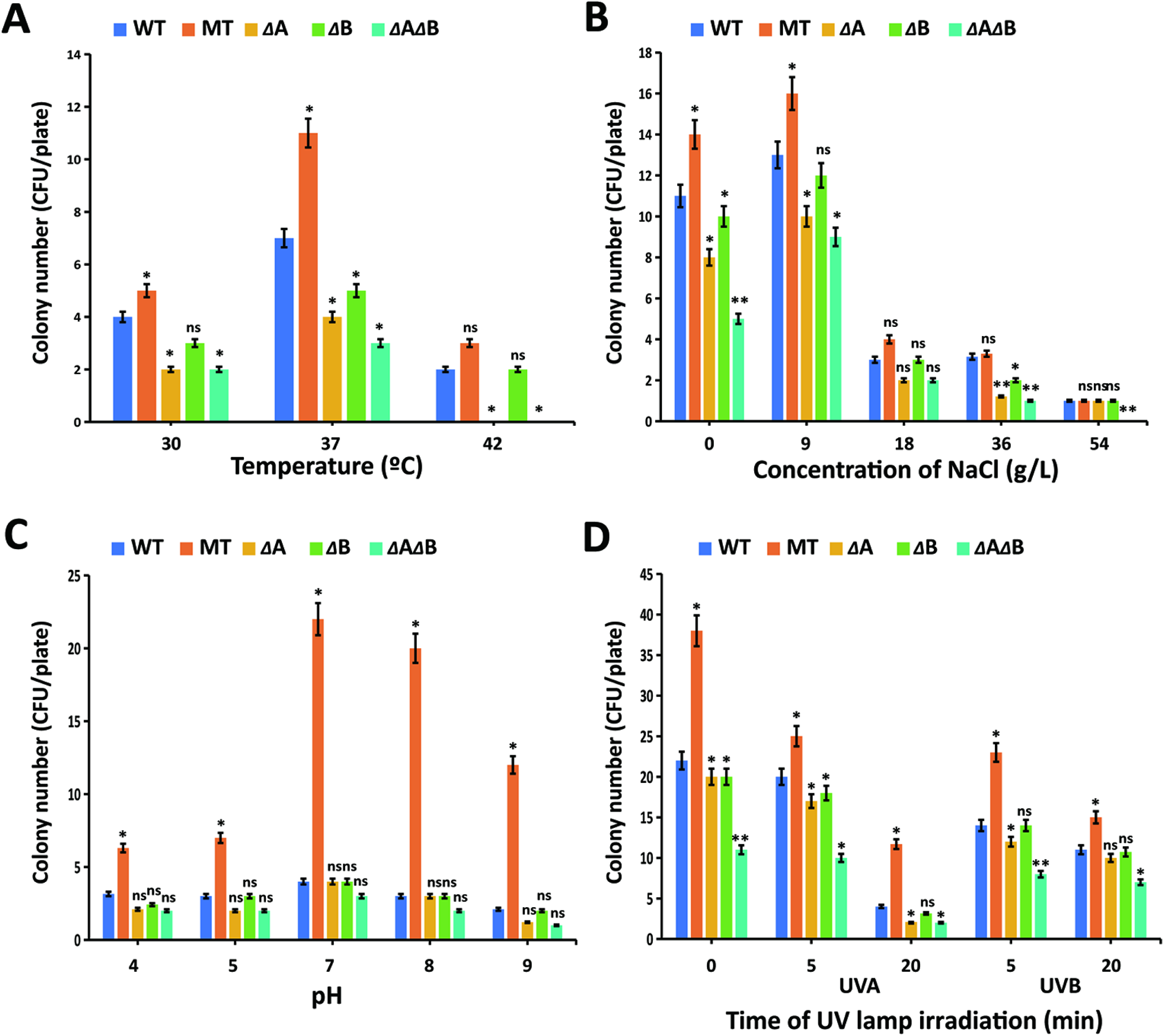
Effects of *spoVF* operon subunits on the tolerance of *Bacillus subtilis* strains to different stress conditions. **(A)** Temperature tolerance. **(B)** NaCl tolerance. **(C)** pH tolerance. **(D)** UV tolerance. *Abbreviations:* WT, wildtype *B. subtilis* strain 168; MT, mutant obtained after promoter replacement (8); *Δ*A, *spoVFA* knockout strain; *Δ*B, *spoVFB* knockout strain; *Δ*A*Δ*B, *spoVFA* and *spoVFB* double knockout strain. Means were submitted to Student’s t test, with significance values at the *p* < 0.05 (*), *p* < 0.01 (**), *p* < 0.001 (***), and *p* < 0.0001 (****) levels (ns = no significance). Means at different culture conditions were compared with those of WT.

To study the tolerance of the bacterial strains to osmotic stress, the strains were grown under different NaCl concentrations, and the survival rates were measured (Fig. 6B). The highest CFU values for all strains were observed when culturing under 9 g/L NaCl, indicating that this is the optimum NaCl concentration for *B. subtilis* growth. As the concentration of NaCl increased, the CFU values of all strains decreased. When 36 g/L NaCl was applied, MT and WT showed similar CFU values, which were higher than those of *Δ*B, *Δ*A, and *Δ*A*Δ*B. The CFU per plate of MT, WT, *Δ*B, *Δ*A, and *Δ*A*Δ*B on 36 g/L NaCl were 3.4 ± 0.5, 3.2 ± 0.5, 2.1 ± 0.3, 1.3 ± 0.1, and 1.1 ± 0.1, respectively. When 54 g/L NaCl was applied, *Δ*A*Δ*B did not grow, whereas the CFU values of the rest of the strains strongly decreased. These results indicated that the expression of the *spoVF* operon is involved the survival ability of *B. subtilis* under osmotic stress, and *spoVFA* appears to play a more important role than *spoVFB*. Interestingly, the higher expression of the *spoVF* operon in MT enhanced the survival rates under high NaCl concentrations.

Under different pH conditions, the number of CFU produced by MT was significantly higher than the other strains (Fig. 6C). This effect could be detected especially under alkaline pH conditions. Under acidic (pH = 4, 5) and alkaline (pH = 9) conditions, the CFU values of *Δ*A and *Δ*A*Δ*B were significantly lower than those of WT, while *Δ*B strain showed similar survival rates comparted to WT. For example, at pH 4, the CFU per plate formed by MT, WT, *Δ*B, *Δ*A, and *Δ*A*Δ*B were 6.2 ± 0.4, 3.4 ± 0.2, 2.3 ± 0.2, 2.2 ± 0.1 and 2.1 ± 0.1, respectively. At pH 9, the CFU per plate formed by MT, WT, *Δ*B, *Δ*A, and *Δ*A*Δ*B were 12.4 ± 0.5, 2.2 ± 0.1, 2.1 ± 0.1, 1.2 ± 0.1, and 1.1 ± 0.1, respectively. These results indicated that *spoVFA* plays a major role in the tolerance of *B. subtilis* to extreme pH conditions.

Both UVA and UVB irradiations decreased the survival rates of the bacterial strains (Fig. 6D). As the irradiation time increased, the survival rates decreased. For example, application of UVA for 5 and 20 min on WT resulted in 20.0 ± 1.1 and 4.2 ± 0.2 CFU per plate, respectively, whereas application of UVB for 5 and 20 min on WT resulted in 14.5 ± 1.6, and 11.4 ± 1.4 CFU per plate, respectively. Generally, the inhibitory effects of UVA irradiation were greater than those of UVB irradiation. MT strain showed the highest survival rates, whereas *Δ*A*Δ*B strain showed the lowest survival rates. When applying UVA for 20 min, the CFU values of MT, WT, *Δ*B, *Δ*A, and *Δ*A*Δ*B were 12.2 ± 0.4, 4.2 ± 0.1, 3.2 ± 0.1, 2.2 ± 0.1, and 2.1 ± 0.1, respectively.

When applying UVB for 20 min, the CFU values of MT, WT, *Δ*B, *Δ*A, and *Δ*A*Δ*B were 15.5 ± 0.5, 11.4 ± 1.4, 11.2 ± 0.3, 10.5 ± 0.2, and 7.2 ± 0.1, respectively.

### Roles of *spoVF* subunits in nutrient competition with *C. fimbriata*

The germination rate of *C. fimbriata* spores increased from 90% in ddH_2_O to 99% and 100% when cultured in 0.5% and 5% sweet potato juice solutions, respectively (Fig. 7). There results can be explained considering that the sweet potato-containing solutions have more nutrients than ddH_2_O, promoting *C. fimbriata* spore germination. MT, WT, *Δ*A, *Δ*B, and *Δ*A*Δ*B reduced *C. fimbriata* germination in ddH_2_O. The highest inhibitory effects in ddH_2_O were detected when using MT and WT, which reduced *C. fimbriata* germination rate by 100%. On the other hand, *Δ*A, *Δ*B, and *Δ*A*Δ*B reduced *C. fimbriata* germination rate in ddH_2_O by 90%, 92%, and 86%, respectively. In 0.5% and 5% sweet potato juice solutions, MT strain allowed the highest inhibitory rates, followed by WT, *Δ*B, *Δ*A, and *Δ*A*Δ*B. MT reduced *C. fimbriata* germination rates in 0.5% and 5% sweet potato juice solutions by 92% and 79%, respectively. In contrast, *Δ*A*Δ*B reduced *C. fimbriata* germination rates in 0.5% and 5% sweet potato juice solutions by only 75% and 47%, respectively.

**FIG 7.**
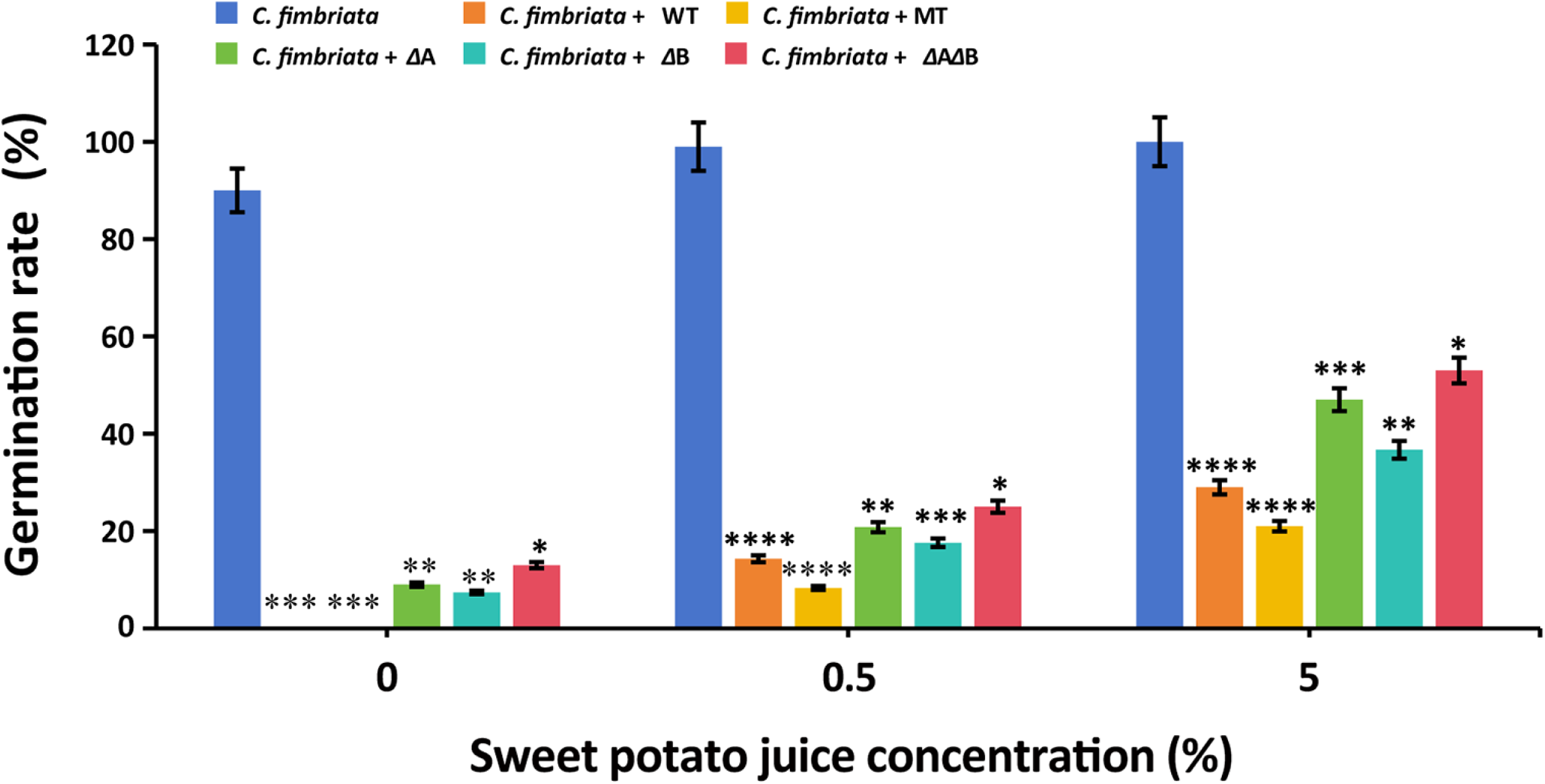
Effects of *spoVF* operon subunits on *Bacillus subtilis* ability to compete for nutrients. This ability was measured according to the inhibitory effects of the bacterial strains on *Ceratocystis fimbriata* conidia germination. The method reported by Di Francesco et al. was used in the assay (44). This method used two polystyrene cylinders separated with a hydrophilic polytetrafluoroethylene (PTFE) membrane, with a pore size of 0.45 μm, which allowed the exchange of nutrients but not cells. *Abbreviations:* WT, wildtype *B. subtilis* strain 168; MT, mutant obtained after promoter replacement (8); *Δ*A, *spoVFA* knockout strain; *Δ*B, *spoVFB* knockout strain; *Δ*A*Δ*B, *spoVFA* and *spoVFB* double knockout strain. Means were submitted to Student’s t test, with significance values at the *p* < 0.05 (*), *p* < 0.01 (**), *p* < 0.001 (***), and *p* < 0.0001 (****) levels (ns = no significance). The control experiment performed with *C. fimbriata* conidia alone, in the absence of bacterial strains. Means at different sweet potato juice concentrations were compared with the control group.

### Roles of *spoVF* subunits in *B. subtilis* in vitro antifungal activity against *C. fimbriata*

When *C. fimbriata* was co-cultured with MT and WT, it showed more obvious biased mycelial growth in comparison to the co-cultures of *C. fimbriata* with the other bacterial strains (Fig. 8). This indicated that the *in vitro* inhibitory activities of MT and WT were higher than those of *Δ*B, *Δ*A, and *Δ*A*Δ*B. Consistently, the fungal cells near MT and WT showed blue colour after staining with Evans blue, and not color after staining with Neutral red, indicating that these cells were death. The number of fungal cells with blue color after inoculation of MT strain was higher than after inoculation of WT. In contrast, the fungal cells far away from WT and MT showed no colour after staining with Evans blue and red color after staining with Neutral red, indicating that these cells were alive. Although *Δ*A and *Δ*B produced small amounts of DPA, the fungal cells near *Δ*A and *Δ*B showed no colour after staining with Evans blue. The same result was obtained when culturing *Δ*A*Δ*B. Instead, the fungal cells near *Δ*A, *Δ*B, and *Δ*A*Δ*B showed red colour after staining with Neutral red, indicating that these cells were alive. These results indicated that the ability of *B. subtilis* 168 to cause *C. fimbriata* cell death is eliminated when knocking out the *spoVF* operon subunits.

**FIG 8.**
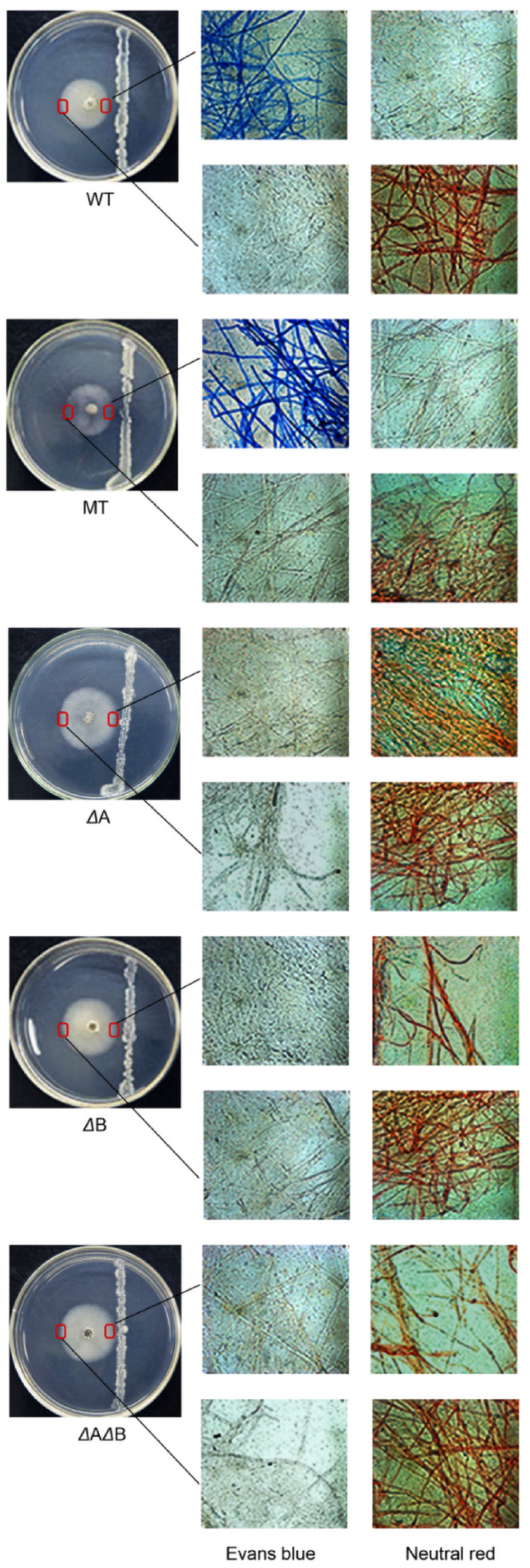
Effects of *spoVF* operon subunits on the *in vitro* antifungal activity of *Bacillus subtilis* against *Ceratocystis fimbriata*. Colour after Evans blue staining indicated that the cells were death, whereas colour after Neutral red staining indicated that the cells were alive. *Abbreviations:* WT, wildtype *B. subtilis* strain 168; MT, mutant obtained after promoter replacement (8); *Δ*A, *spoVFA* knockout strain; *Δ*B, *spoVFB* knockout strain; *Δ*A*Δ*B, *spoVFA* and *spoVFB* double knockout strain.

### Roles of *spoVF* subunits in *B. subtilis* efficacy against *C. fimbriata* on sweet potato

The inhibitory efficacy of the bacterial strains was measured according to their ability to reduce sweet potato black rot disease incidence and lesion length. As it can be observed in Table 2, the five strains reduced the symptoms caused by *C. fimbriata* on sweet potato. The higher the concentration of the strains, the higher the inhibitory efficacy. At the same cell concentration, MT strain was the most efficient, followed by WT, *Δ*B, *Δ*A, and *Δ*A*Δ*B. When comparing with the control experiment, which was carried out in the absence of bacterial strains, the lesion diameter decreased by 74% and 91% after applying 1 × 10^7^ and 1 × 10^8^ CFU/mL MT, respectively (Fig. 9). However, the lesion diameter only decreased by 12% and 43% when using 1 × 10^7^ and 1 × 10^8^ CFU/mL *Δ*A*Δ*B, respectively. The disease incidence decreased by 62% and 92% when using 1 × 10^7^ and 1 × 10^8^ CFU/mL MT, respectively, whereas the disease incidence decreased by only 12% and 32% when applying 1 × 10^7^ and 1 × 10^8^ CFU/mL *Δ*A*Δ*B, respectively.

**FIG 9.**
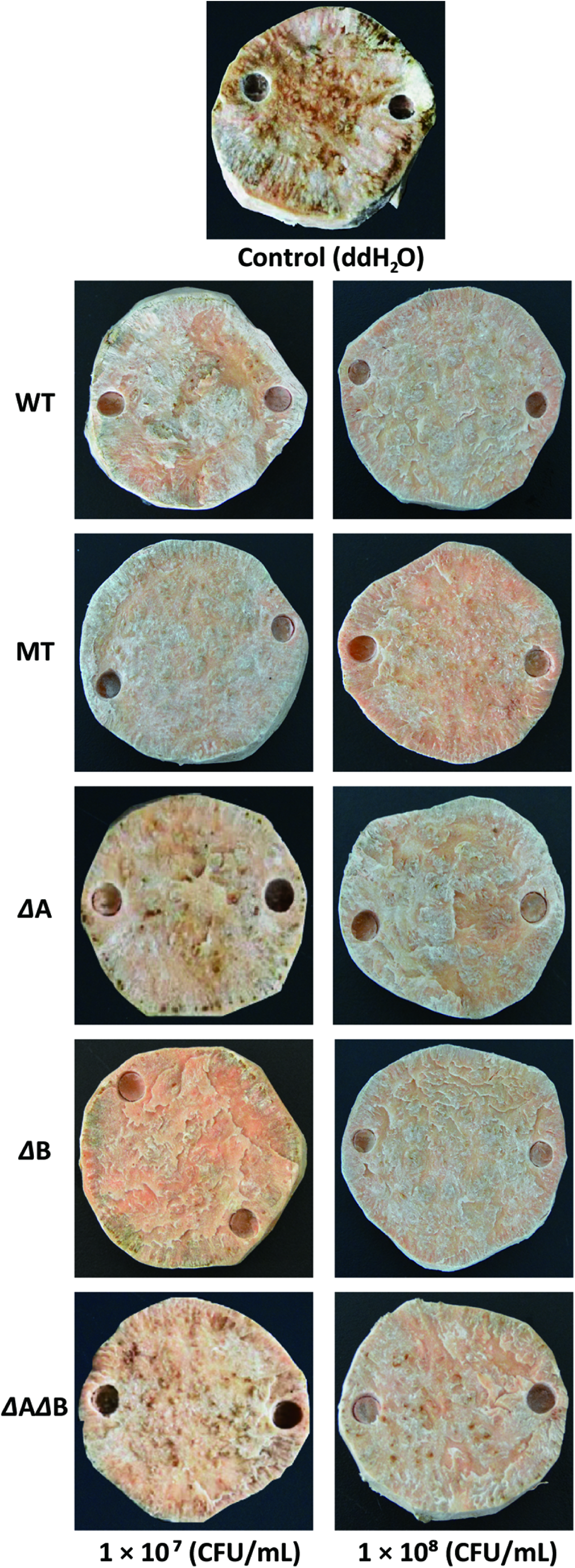
Effects of *spoVF* operon subunits on the biocontrol ability of *Bacillus subtilis* against *Ceratocystis fimbriata* on sweet potato. Sterilized ddH_2_O was sprayed in the control experiment. **(A)** Photos showing the lesions caused by *C. fimbriata* on sweet potato slices. **(B)** Lesion length caused by *C. fimbriata* on sweet potato slices. *Abbreviations:* WT, wildtype *B. subtilis* strain 168; MT, mutant obtained after promoter replacement (8); *Δ*A, *spoVFA* knockout strain; *Δ*B, *spoVFB* knockout strain; *Δ*A*Δ*B, *spoVFA* and *spoVFB* double knockout strain.

**TABLE 2.** Lesion length and disease incidence of *Ceratocystis fimbriata* causing black rot disease after treatment with *Bacillus subtilis* strains.

| Antifungal agent | Cell concentration (CFU/mL) | Disease incidence (%)<sup>a</sup> | Lesion length (cm) | Significance<sup>b</sup> |
| --- | --- | --- | --- | --- |
| Control<sup>c</sup> | - | 82.5 | 3.40 ± 0.35 | - |
| WT | 1 × 10<sup>7</sup> | 53.8 | 1.29 ± 0.25 | * |
|  | 1 × 10<sup>8</sup> | 32.5 | 0.41 ± 0.23 | * |
| MT | 1 × 10<sup>7</sup> | 31.3 | 0.90 ± 0.21 | * |
|  | 1 × 10<sup>8</sup> | 6.3 | 0.31 ± 0.14 | * |
| ΔA | 1 × 10<sup>7</sup> | 65.0 | 2.79 ± 0.34 | * |
|  | 1 × 10<sup>8</sup> | 42.5 | 1.83 ± 0.39 | * |
| ΔB | 1 × 10<sup>7</sup> | 61.3 | 1.54 ± 0.19 | * |
|  | 1 × 10<sup>8</sup> | 36.3 | 0.94 ± 0.10 | * |
| ΔAΔB | 1 × 10<sup>7</sup> | 72.5 | 3.00 ± 0.38 | * |
|  | 1 × 10<sup>8</sup> | 56.3 | 1.95 ± 0.37 | * |

<sup>a</sup>The infection rate is the ratio between the number of infections and the total number of tests.
<sup>b</sup>Significance level at $p < 0.05$ (\*). The treatment groups were compared with the control group.
<sup>c</sup>Control experiment was performed by spraying water.

## DISCUSSION

This study revealed that the knockout of *spoVFA* and *spoVFB* genes has a significant impact on *B. subtilis* cell morphology, with *spoVFA* having a more significant effect than *spoVFB*. This suggests that the *spoVF* operon may have a regulatory effect on genes related to cell wall metabolism. In agreement with our results, changes in DPA content in *B. subtilis* spores were reported to increase hydration, altering the normal cell morphology (49). Zhang et al. reported that the *spoVF* operon has complex regulatory relationships with some genes, including genes related to cell wall metabolism, such as peptidoglycan hydrolase related genes, whose expression is strictly regulated to ensure the correct synthesis and remodeling of the cell wall during spore formation (50). Dersch et al. found that MreB is an ortholog of *B. subtilis* actin, which aggregates into filamentous structures at the cell periphery and forms complexes with proteins related to cell wall synthesis, thereby affecting cell morphology and mechanical rigidity (51). One of the functions of MreB protein is to ensure the uniform distribution of new peptidoglycan insertion sites, which is a necessary condition for bacteria to maintain rod shape (52). The MreB protein can recruit enzymes related to the synthesis of peptidoglycans and phospholipids, and participate in the maintenance of cell morphology by regulating the bacterial cell wall synthesis pathway. When the MreB of rod-shaped bacteria, such as *Escherichia coli* and *Salmonella* sp., was knocked out, the cell shape changed from rod to spherical (53). Bratton et al. found that the inhibition of MreB expression in *E. coli* can lead to decreased cellular mechanical rigidity, and cells lacking MreB loosed their rod morphology (54). In areement with those previous studies, after knocking out *spoVFA* and *spoVFB* separately or simultaneously in *B. subtilis*, the expression levels of *merB1* and *merB2* were significantly reduced, and the cell morphology changed from rod to spherical. Therefore, we speculate that the *spoVF* operon may affect cell wall synthesis by influencing peptidoglycan synthesis, thereby affecting cell morphology.

During DPA biosynthesis, cofactors ATP, NADPH, and NAD^+^ are consumed and not regenerated (38). The expression of the *spoVF* operon is enhanced in vegetative cells in MT strain, resulting in enhanced DPA synthesis compared to WT. If we consider MT as template, the overproduction of DPA may result in an increase of ATP, NADPH, and NAD^+^ demands (Fig. 10). The pentose phosphate pathway (PPP) is related to biomass synthesis and is the main pathway for generating NADPH (55). Cell growth rate and NADPH content induction in other microorganisms has been reported to be a response related to NADPH demand (56).

**FIG 10.**
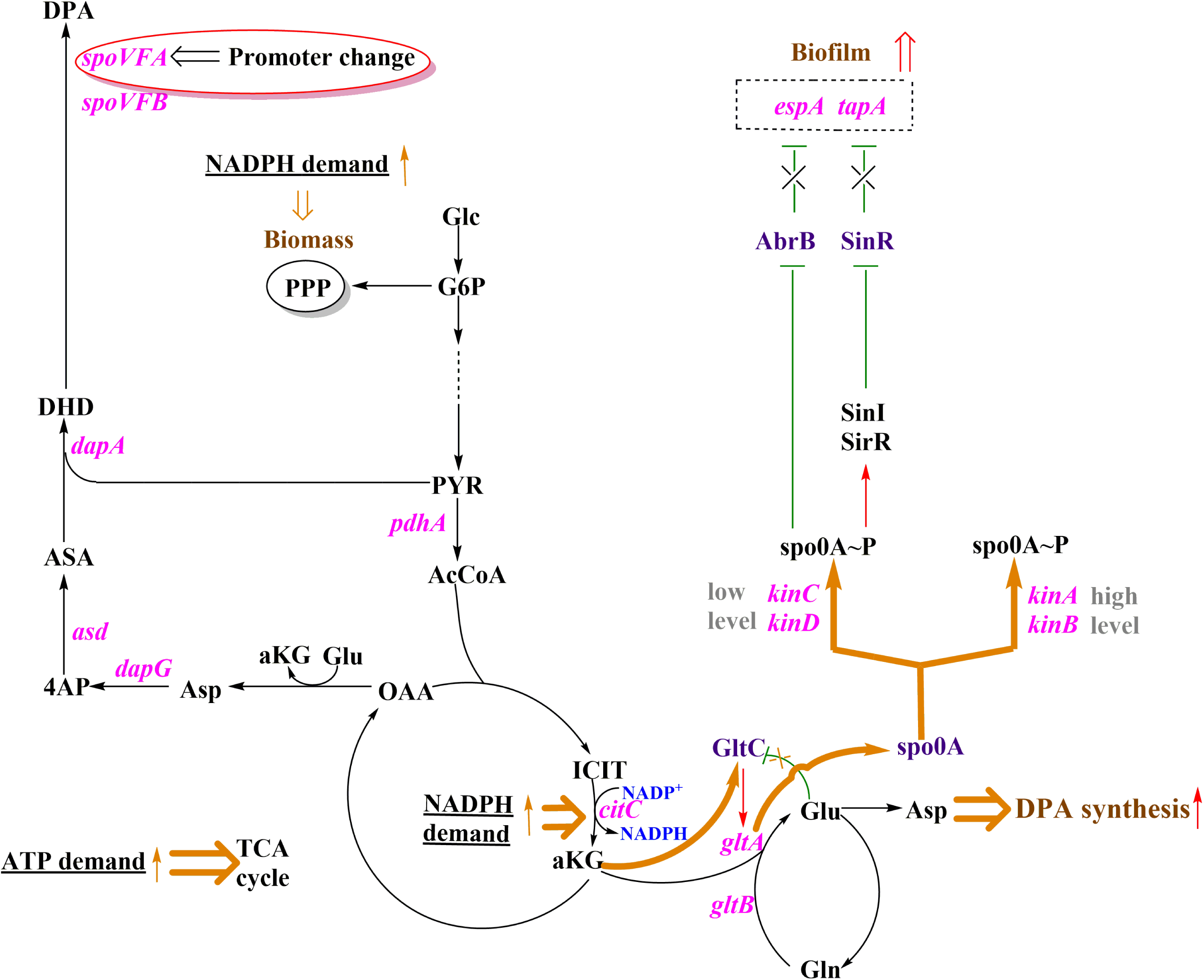
Proposed regulatory mechanism between the *spoVF* operon and biofilm formation by *Bacillus subtilis*. This scheme aims to explain how biofilm formation is regulated in the MT strain. *Abbreviations:* MT, mutant obtained after promoter replacement (8).

Following with MT, the increase in ATP demand may induce an increase in TCA cycle flux, as well as an increase in NADPH demand. This may in turn promote isocitrate dehydrogenase-catalyzed conversion from isocitrate into α-ketoglutarate, which is in agreement with the higher expression level of *citC* (Fig. 10). The accumulation of α-ketoglutarate may induce the expression of transcription factor GltC, thereby activating the transcription of GOGAT-encoding gene *gltA* (57). GOGAT catalyzes the transformation of α-ketoglutaric acid into glutamic acid. Thus, glutamic acid may be used as a precursor for DPA synthesis, enhancing DPA accumulation. The expression and phosphorylation level of transcription factor spo0A was previously reported to decrease in the *ΔgltA* mutant (58). Consistently, the upregulation of *gltA* expression induced the expression of spo0A in this study.

The main component of *B. subtilis* biofilm is the extracellular polysaccharide TasA amyloid fiber, which biosynthetic steps are encoded in the *epsA* and *tapA* operons. The genes in the *epsA* and *tapA* operons are inhibited by the inhibitory transcription factors AbrB and SinR (59). Spo0A is phosphorylated to give spo0A∼P, and the levels of Spo0A phosphorylation can be low, which involves the participation of *kinC* and *kinD* genes, or high, which involves the participation of *kinA* and *kinB* genes (60). The low phosphorylation level of spo0A is a necessary condition for initiating biofilm formation. Low phosphorylation level spo0A∼P can inhibit the transcription of AbrB and induce the expression of SinI and SlrR proteins, thereby inhibiting SinR and ultimately stimulating biofilm formation (Fig. 10). Thus, we can explain why MT produces higher biofilm amounts and establish the network that may interconnect the *spoVF* operon with the biofilm inhibitory factors AbrB and SinR.

In contrast, when the *spoVFA* and *spoVFB* subunits are knocked-out, DPA synthesis must be weakened, leading to decreased demands for ATP and NADPH.

This may in turn reduce the cell growth rate and NADPH content. In consequence, the decreased ATP requirements may lead to decreased TCA cycle flux, reducing isocitrate dehydrogenase-catalyzed formation of α-ketoglutaric acid. The inhibition of α-ketoglutaric acid formation may inhibit the expression of the transcriptional factor GltC, downregulating *gltA* (57). This may in turn lead to a decrease in spo0A expression and phosphorylation. Without the activity of spo0A∼P, the transcription suppressors AbrB and SinR may be activated, inhibiting the expression of *epsA* and *tapA* and subsequently biofilm formation in *Δ*A, *Δ*B, and *Δ*A*Δ*B strains.

The lower tolerance of *Δ*A, *Δ*B, and *Δ*A*Δ*B strains compared to the WT strain can be explained considering that PDA levels are much lower in *Δ*A and *Δ*B strains, and *Δ*A*Δ*B strain can not produce DPA. As indicated in the Introduction section, DPA is involved in *B. subtilis* tolerance to heat, UV radiation, dryness, and extreme pH stresses (32,33). These effects have been related to the key role of DPA in the dehydration of spores, which seems to increase their tolerance to harsh environmental conditions (61). Release of DPA from *B. subtilis* spores to the medium results in decreased resistance to pressure and metals (62,63).

Competition for nutrients in *Bacillus* is one of the most important antifungal mechanisms (64–66). The higher efficacies of the bacterial strains to inhibit *C. fimbriata* germination in ddH_2_O than in 0.5% and 5% sweet potato juice solutions can be explained considering that there is a greater availability of nutrients for the pathogen in the sweet potato juice solutions and, for this reason, nutrient deprivation effects are not such obvious. Interestingly, our results indicated that the *spoVF* subunits play a key role in the ability of *B. subtilis* to compete for nutrients with fungal pathogens. The higher inhibitory activities observed with WT and MT than with *Δ*A, *Δ*B and *Δ*A*Δ*B can be explained considering the faster growth and glucose consumption of WT and MT compared with the other bacterial strains. This may result in turn in a higher deprivation of nutrients when using WT and MT than when using the other bacterial strains.

Given that DPA is one of the main antifungal metabolites from *B. subtilis* 168, it is easy to explain the lower inhibitory activities of *Δ*A, *Δ*B, and *Δ*A*Δ*B strains *in vitro* and *in vivo* compared to MT and WT. In addition, *Δ*A, *Δ*B, and *Δ*A*Δ*B strains showed less biofilm formation, and subsequently, lower attachment capacity in the sweet potato tubers. And the decrease in growth rate affected negatively the nutrient competition ability of *Δ*A, *Δ*B, and *Δ*A*Δ*B strains. All these alterations in turn limited the antifungal efficacies of *Δ*A, *Δ*B, and *Δ*A*Δ*B strains for the management of black rot disease in sweet potatoes. The obtained results also offer a new understanding regarding how to develop efficient biocontrol strategies for the management of *C. fimbriata*. It must be noted that alternative methods for the control of this hazardous pathogen are lacking (67–69).

## Conclusion

In summary, the role of the *spoVF* operon subunits in biofilm formation and cell morphology regulation was confirmed, and the mechanism of *spoVF* operon regulating biofilm formation was proposed in this study. *Δ*A, *Δ*B, and *Δ*A*Δ*B strains showed altered morphology and decreased growth rates, glucose consumption, DPA production, biofilm formation, and stress tolerance to heat, slat, extreme pH conditions, and UV irradiation compared to the WT strain. The decrease in biofilm formation resulted in lower ability to attach to sweet potato surface. Additionally, *Δ*A, *Δ*B, and *Δ*A*Δ*B strains showed lower ability to compete for nutrients and decreased antifungal activity *in vitro* and *in vitro* against *C. fimbriata*. The interconnection between the simultaneous inhibition of the *spoVF* operon and biofilm formation appears to be related to a decrease in the TCA cycle flux, which in turn activates biofilm inhibitory transcription factors AbrB and SinR. In contrast, MT strain, which had increased DPA production via replacement of the *spoVF* promoter, showed the opposite effects and inhibited *C. fimbriata* in *in vitro* and *in vitro* assays to a greater extent than the WT strain. This study demonstrated that both *spoVF* operon subunits are involved in the biocontrol properties of *B. subtilis*, with the *spoVFA* subunit playing a more important role than the *spoVFB* subunit. This study reveals relevant insights to understand the metabolic networks involved in the antifungal ability of the major biocontrol agent *B. subtilis*. The results obtained in this study may help to develop more efficient biocontrol products based on *B. subtilis*.

## ACKNOWLEDGEMENTS

This research was funded by the National Natural Science Foundation of China (32302433 and 32172441), the Large Instruments Open Foundation of Nantong University (KFJN2440, KFJN2425, and KFJN2480), the College Student Innovation Training Program Project (202310304028Z), and the Postgraduate Research & Practice Innovation Program of Jiangsu Province (KYCX24_3615).

## DATA AVAILABILITY

The strains used in this study will be made available upon request.

## ADDITIONAL FILES

The following material is available online.

Supplemental Material

Tables S1 to S3.

## REFERENCES

1 Pomerleau M, Charron-Lamoureux V, Léonard L, Grenier F, Rodrigue S, Beauregard PB. 2024. Adaptive laboratory evolution reveals regulators involved in repressing biofilm development as key players in *Bacillus subtilis* root colonization. Msystems 9:e00843–23. 10.1128/msystems.00843-23

2 Cavaglieri L, Orlando J, Rodríguez M, Chulze S, Etcheverry M. 2005. Biocontrol of *Bacillus subtilis* against *Fusarium verticillioides in vitro* and at the maize root level. Res Microbiol 156:748–754. 10.1016/j.resmic.2005.03.001

3 Fan HY, Zhang ZW, Li Y, Zhang X, Duan YM, Wang Q. 2017. Biocontrol of bacterial fruit blotch by *Bacillus subtilis* 9407 via surfactin-mediated antibacterial activity and colonization. Front Microbiol 8:1973. 10.3389/fmicb.2017.01973

4 Munir S, Li YM, He PF, He PB, He PJ, Cui WY, Wu YX, Li XY, He YQ. 2018. *Bacillus subtilis* L1-21 possible assessment of inhibitory mechanism against phytopathogens and colonization in different plant hosts. Pak J Agr Sci 55:996– 1002. 10.21162/PAKJAS/18.7750

5 Dunlap CA, Bowman MJ, Rooney AP. 2019. Iturinic lipopeptide diversity in the *Bacillus subtilis* species group-important antifungals for plant disease biocontrol applications. Front Microbiol 10:1794. 10.3389/fmicb.2019.01794

6 Penha RO, Vandenberghe LPS, Faulds C, Soccol VT, Soccol CR. 2020. *Bacillus* lipopeptides as powerful pest control agents for a more sustainable and healthy agriculture: recent studies and innovations. Planta 251:70. 10.1007/s00425-020-03357-7

7 Song XG, Han MH, He F, Wang SY, Li HC, Wu GC, Huang ZG, Liu D, Liu FQ, Laborda P, Shi XC. 2020. Antifungal mechanism of dipicolinic acid and its efficacy for the biocontrol of pear Valsa canker. Front Microbiol 11:958. 10.3389/fmicb.2020.00958

8 Wang T, Wang XC, Han MH, Song XG, Yang DJ, Wang SY, Laborda P, Shi XC. 2021. Enhanced *spoVF* operon increases host attachment and biocontrol ability of *Bacillus subtilis* for the management of *Ceratocystis fimbriata* in sweet potato. Biol Control 161:104651. 10.1016/j.biocontrol.2021.104651

9 Zhang D, Yu SQ, Yang YQ, Zhang JL, Zhao DM, Pan Y, Fan SS, Yang ZH, Zhu JH. 2020. Antifungal effects of volatiles produced by *Bacillus subtilis* against *Alternaria solani* in potato. Front Microbiol 11:1196. 10.3389/fmicb.2020.01196

10 Dobrange E, Porras-Dominguez JR, van den Ende W. 2024. The complex GH32 enzyme orchestra from *Priestia megaterium* holds the key to better discriminate sucrose-6-phosphate hydrolases from other β-fructofuranosidases in bacteria. J Agric Food Chem 72:1302–1320. 10.1021/acs.jafc.3c06874

11 Yi YJ, Yin YN, Yang YA, Liang YQ, Shan YT, Zhang CF, Zhang YR, Liang ZP. 2022. Antagonistic activity and mechanism of *Bacillus subtilis* XZ16-1 suppression of wheat powdery mildew and growth promotion of wheat. Phytopathology 112:2476–2485. 10.1094/phyto-04-22-0118-r

12 Eom JS, Lee SY, Choi HS. 2014. *Bacillus subtilis* HJ18-4 from traditional fermented soybean food inhibits *Bacillus cereus* growth and toxin-related genes. J Food Sci 79:M2279–87. 10.1111/1750-3841.12569

13 Radzhabov RU, Davranov K. 2010. Metabolites of *Bacillus subtilis* SKB 256, growth inhibitors of phytopathogenic fungi. Chem Nat Compd 46:160–162. 10.1007/s10600-010-9556-y

14 Chen Y, Zhou YD, Laborda P, Wang HL, Wang R, Chen X, Liu FQ, Yang DJ, Wang SY, Shi XC, Laborda P. 2021. Mode of action and efficacy of quinolinic acid for the control of *Ceratocystis fimbriata* on sweet potato. Pest Manag Sci 77:4564–4571. 10.1002/ps.6495

15 Cong H, Sun Y, Li CG, Zhang YJ, Wang YM, Ma DF, Jiang JH, Li LW, Li LD. 2024. The APSES transcription factor CfSwi6 is required for growth, cell wall integrity, and pathogenicity of *Ceratocystis fimbriata*. Microbiol Res 281:127624. 10.1016/j.micres.2024.127624

16 Liu M, Meng QC, Wang S, Yang KL, Tian J. 2023. Research progress on postharvest sweet potato spoilage fungi *Ceratocystis fimbriata* and control measures. Food Biosci 53:102627. 10.1016/j.fbio.2023.102627

17 Stahr M, Quesada-Ocampo LM. 2021. Effects of water temperature, inoculum concentration and age, and sanitizers on infection of *Ceratocystis fimbriata*, causal agent of black rot in sweet potato. Plant Dis 105:1365–1372. 10.1094/pdis-07-20-1475-re

18 Latif MZ. 2023. Morphology, pathogenicity and physiology of *Ceratocystis fimbriata* causing black rot disease of *Colocasia esculenta*. Pak J Agr Sci 60:265 –272. 10.21162/pakjas/23.635

19 Herrera-Balandrano DD, Wang SY, Wang B, Yang DJ, Shi XC, Laborda P. 2023. Methods for the control of the soil-borne pathogen *Ceratocystis fimbriata* on sweet potato: A mini review. Pedosphere in press. 10.1016/j.pedsph.2023.12.009

20 Mohsin SM, Hasanuzzaman M, Parvin K, Morokuma M, Fujita M. 2021. Effect of tebuconazole and trifloxystrobin on *Ceratocystis fimbriata* to control black rot of sweet potato: processes of reactive oxygen species generation and antioxidant defense responses. World J Microbiol Biotechnol 37:148. 10.1007/s11274-021-03111-5

21 Scruggs AC, Basaiah T, Adams ML, Quesada-Ocampo LM. 2017. Genetic diversity, fungicide sensitivity, and host resistance to *Ceratocystis fimbriata* infecting sweet potato in North Carolina. Plant Dis 101:994–1001. 10.1094/pdis-11-16-1583-re

22 Jiang YH, Shi XC, Wu T, Du H, Pang YB, Zhou R, Yin HP, Herrera-Balandrano DD, Yang DJ, Lu AM, Laborda P, Polo V, Wang SY. 2024. Synthesis and antifungal activity of novel amide derivatives from quinic acid against the sweet potato pathogen *Ceratocystis fimbriata*. Pest Manag Sci in press. 10.1002/ps.8527

23 Oleńska E, Małek W, Wójcik M, Swiecicka L, Thijs S, Vangronsveld J. 2020. Beneficial features of plant growth-promoting rhizobacteria for improving plant growth and health in challenging conditions: a methodical review. Sci Total Environ 743:140682. 10.1016/j.scitotenv.2020.140682

24 Tsiantas P, Tzanetou EN, Karasali H, Kasiotis KM. 2021. A dieldrin case study: Another evidence of an obsolete substance in the European soil environment. Agriculture 11:314. 10.3390/agriculture11040314

25 Xu MJ, Guo JH, Li TJ, Zhang CM, Peng X, Xing K, Qin S. 2021. Antibiotic effects of volatiles produced by *Bacillus tequilensis* XK29 against the black spot disease caused by *Ceratocystis fimbriata* in postharvest sweet potato. J Agric Food Chem 69:13045–13054. 10.1021/acs.jafc.1c04585

26 Ru YR, Liu JW, Xu PJ, Gao WH, Sun D, Zhu JR, Liu C, Liu WJ. 2022. Application of the biosurfactant produced by *Bacillus velezensis* MMB-51 as an efficient synergist of sweet potato foliar fertilizer. J Surfactants Deterg 25:743– 756. 10.1002/jsde.12610

27 Jiang LM, Jeong JC, Lee JS, Park J, Yang JW, Lee MH, Choi SH, Kim DH, Kim SW, Lee J. 2019. Potential of *Pantoea dispersa* as an effective biocontrol agent for black rot in sweet potato. Sci Rep 9:16354. 10.1038/s41598-019-52804-3

28 Zhang Y, Li TJ, Liu YF, Li XY, Zhang CM, Feng ZZ, Peng X, Li ZY, Qin S, Xing K. 2019. Volatile organic compounds produced by *Pseudomonas chlororaphis* subsp. *aureofaciens* SPS-41 as biological fumigants to control *Ceratocystis fimbriata* in postharvest sweet potatoes. J Agric Food Chem 67:3702–3710. 10.1021/acs.jafc.9b00289

29 Hong CE, Jeong H, Jo SH, Jeong JC, Kwon SY, An D, Park JM. 2016. A leaf-inhabiting endophytic bacterium, *Rhodococcus* sp. KB6, enhances sweet potato resistance to black rot disease caused by *Ceratocystis fimbriata*. J Microbiol Biotechnol 26:488–492. 10.4014/jmb.1511.11039

30 Li X, Li B, Cai S, Zhang Y, Xu M, Zhang C, Yuan B, Xing K, Qin S. 2020. Identification of rhizospheric actinomycete *Streptomyces lavendulae* SPS-33 and the inhibitory effect of its volatile organic compounds against *Ceratocystis Fimbriata* in postharvest sweet potato (*Ipomoea Batatas* (L.) Lam.). Microorganisms 8:319. 10.3390/microorganisms8030319

31 Gong Y, Liu JQ, Xu MJ, Zhang CM, Gao J, Li CG, Xing K, Qin S. 2022. Antifungal volatile organic compounds from *Streptomyces setonii* WY228 control black spot disease of sweet potato. Appl Environ Microbiol 88:e0231721. 10.1128/aem.02317-21

32 Ramírez-Guadiana FH, Meeske AJ, Rodrigues CDA, Barajas-Ornelas RD, Kruse AC, Rudner DZ. 2017. A two-step transport pathway allows the mother cell to nurture the developing spore in *Bacillus subtilis*. Plos Genet 13:e1007015. 10.1371/journal.pgen.1007015

33 Takahashi F, Sumitomo N, Hagihara H, Ozaki K. 2015. Increased dipicolinic acid production with an enhanced *spoVF* operon in *Bacillus subtilis* and medium optimization. Biosci Biotechnol Biochem 79:505–511. 10.1080/09168451.2014.978261

34 Orsburn BC, Melville SB, Popham DL. 2010. EtfA catalyses the formation of dipicolinic acid in *Clostridium perfringens*. Mol Microbiol 75:178–186. 10.1111/j.1365-2958.2009.06975.x

35 Wen J, Vischer NOE, de Vos AL, Manders EMM, Setlow P, Brul S. 2022. Organization and dynamics of the SpoVAEa protein and its surrounding inner membrane lipids, upon germination of *Bacillus subtilis* spores. Sci Rep 12:4944. 10.1038/s41598-022-09147-3

36 Ji XY, Wang BY, Zhang YF, Zhang YJ, Lai YJ, Yang Y, Wang XC, Wang SY, Laborda P, Shi XC. 2024. Dipicolinic acid reduces *Epicoccum sorghinum* symptoms on maize and inhibits tenuazonic acid biosynthesis. Pest Manag Sci 80:6545–6554. 10.1002/ps.8393

37 Sarwar A, Brader G, Corretto E, Aleti G, Ullah MA, Sessitsch A, Hafeez FY. 2018. Qualitative analysis of biosurfactants from *Bacillus* species exhibiting antifungal activity. Plos One 13:e0198107. 10.1371/journal.pone.0198107

38 Toya Y, Hirasawa T, Ishikawa S, Chumsakul O, Morimoto T, Liu S, Masuda K, Kageyama Y, Ozaki K, Ogasawara N, Shimizu H. 2015. Enhanced dipicolinic acid production during the stationary phase in *Bacillus subtilis* by blocking acetoin synthesis. Biosci Biotechnol Biochem 79:2073–2080. 10.1080/09168451.2015.1060843

39 Timmusk S, Behers L, Muthoni J, Muraya A, Aronsson AC. 2017. Perspectives and challenges of microbial application for crop improvement. Front Plant Sci 8:49. 10.3389/fpls.2017.00049

40 Brück HL, Delvigne F, Dhulster P, Jacques P, Coutte F. 2019. Molecular strategies for adapting *Bacillus subtilis* 168 biosurfactant production to biofilm cultivation mode. Bioresour Technol 293:122090. 10.1016/j.biortech.2019.122090

41 Zhang YJ, Herrera-Balandrano DD, Shi XC, Wang SY, Laborda P. 2022. Biocontrol of *Colletotrichum brevisporum* in soybean using a new genistein-producing *Paecilomyces* strain. Biol Control 169:104877. 10.1016/j.biocontrol.2022.104877

42 Xia ML, Wang L, Yang ZX, Chen HZ. 2015. A novel digital color analysis method for rapid glucose detection. Anal Methods 7:6654–6663. 10.1039/C5AY01233C

43 Calvio C, Celandroni F, Ghelardi E, Amati G, Salvetti S, Ceciliani F, Galizzi A, Senesi S. 2005. Swarming differentiation and swimming motility in *Bacillus subtilis* are controlled by *swrA*, a newly identified dicistronic operon. J Bacteriol 187:5356–5366. 10.1128/jb.187.15.5356-5366.2005

44 Sun JL, Zhou CH, Zhao Y, Zhang XF, Chen WY, Zhou Q, Hu B, Gao DM, Raatz L, Wang ZF, Nelson PJ, Jiang YC, Ren N, Bruns CJ, Zhou HJ. 2021. Quiescin sulfhydryl oxidase 1 promotes sorafenib-induced ferroptosis in hepatocellular carcinoma by driving EGFR endosomal trafficking and inhibiting NRF2 activation. Redox Biol 41:101942. 10.1016/j.redox.2021.101942

45 Di Francesco A, Ugolini L, D’Aquino S, Pagnotta E, Mari M. 2017. Biocontrol of *Monilinia laxa* by *Aureobasidium pullulans* strains: Insights on competition for nutrients and space. Int J Food Microbiol 248:32–38. 10.1016/j.ijfoodmicro.2017.02.007

46 Lima BC, Cruz TR, Ribas AF, Santos TB, Cacefo V, Araujo FF. 2023. Gene expression and biochemical profiling in the mitigation of heat stress in common bean using *Bacillus subtilis*. Biol Plantarum 63:213–223. 10.32615/bp.2023.022

47 Wang FR, Deng HY, Wu Q, Sun HJ, Zhang J, Li ZY, Zhang LM, Liu MY. 2024. Biocontrol of black rot of sweetpotato by *Pichia pastoris* recombinant strain expressing chitinase IbChiA. Sci Hortic 329:112979. 10.1016/j.scienta.2024.112979

48 Muhammad MH, Idris AL, Fan X, Guo Y, Yu Y, Jin X, Qiu J, Guan X, Huang T. 2020. Beyond risk: bacterial biofilms and their regulating approaches. Front Microbiol 11:928. 10.3389/fmicb.2020.00928

49 Paidhungat M, Setlow B, Driks A, Setlow P. 2000. Characterization of spores of *Bacillus subtilis* which lack dipicolinic acid. J Bacteriol 182:5505–5512. 10.1128/JB.182.19.5505-5512.2000

50 Zhang JF, Li XY, Li XY, Chen XJ, Niu X, Ren SD, Lu FP. 2021. Advance in the effect of reduction of the *Bacillus subtilis* genome on the expression of heterologous enzymes. Wei Sheng Wu Xue Tong Bao 48:859–872. 10.13344/j.microbiol.china.200403

51 Dersch S, Reimold C, Stoll J, Breddermann H, Heimerl T, Soufo HJD, Graumann PL. 2020. Polymerization of *Bacillus subtilis* MreB on a lipid membrane reveals lateral co-polymerization of MreB paralogs and strong effects of cations on filament formation. BMC Mol Cell Biol 21:76. 10.1186/s12860-020-00319-5

52 van Teeffelen S, Wang SY, Furchtgott L, Huang KC, Wingreen NS, Shaevitz JW, Gitai Z. 2011. The bacterial actin MreB rotates, and rotation depends on cell-wall assembly. Proc Natl Acad Sci U.S.A 108:15822–15827. 10.1073/pnas.1108999108

53 Feng LL, Wang ZW. 2021. Development of morphology engineering for production of bio-based chemicals. Zhongguo Sheng Wu Gong Cheng Za Zhi 37:2211–2222. 10.13345/j.cjb.200516

54 Bratton BP, Shaevitz JW, Gitai Z, Morgenstein RM. 2018. MreB polymers and curvature localization are enhanced by RodZ and predict *E. coli’*s cylindrical uniformity. Nat Commun 9:2797. 10.1038/s41467-018-05186-5

55 Masi A, Mach RL, Mach-Aigner AR. 2021. The pentose phosphate pathway in industrially relevant fungi: crucial insights for bioprocessing. Appl Microbiol Biotechnol 105:4017–4031. 10.1007/s00253-021-11314-x

56 Shi XC, Zou YN, Chen Y, Zheng C, Li BB, Xu JH, Shen XN, Ying HJ. 2016. A water-forming NADH oxidase regulates metabolism in anaerobic fermentation. Biotechnol Biofuels 9:103. 10.1186/s13068-016-0517-y

57 Picossi S, Belitsky BR, Sonenshein AL. 2007. Molecular mechanism of the regulation of *Bacillus subtilis gltAB* expression by GltC. J Mol Biol 365:1298– 1313. 10.1016/j.jmb.2006.10.100

58 Kimura T, Kobayashi K. 2020. Role of glutamate synthase in biofilm formation by *Bacillus subtilis*. J Bacteriol 202:e00120–20. 10.1128/JB.00120-20

59 Chu F, Kearns DB, Branda SS, Kolter R, Losick R. 2006. Targets of the master regulator of biofilm formation in *Bacillus subtilis*. Mol Microbiol 59:1216–1228. 10.1111/j.1365-2958.2005.05019.x

60 Tran TNT, Nguyen TDP, Dinh HT, Bui TT, Ho LH, Nguyen-Phan TX, Khoo KS, Chew KW, Show PL. 2022. Characterization of bacteria type strain *Bacillus*. spp isolated from extracellular polymeric substance harvested in seafood wastewater. J Chem Technol Biot 97:501–508. 10.1002/jctb.6870

61 Khanal S, Kim TD, Begyn K, Duverger W, Kramer G, Brul S, Rajkovic A, Devlieghere F, Heyndrickx M, Schymkowitz J, Rousseau F, Broussolle V, Michiels C, Aertsen A. 2024. Mechanistic insights into the adaptive evolvability of spore heat resistance in *Bacillus cereu*s sensu lato. Int J Food Microbiol 418:110709. 10.1016/j.ijfoodmicro.2024.110709

62 Margosch D, Gänzle MG, Ehrmann MA, Vogel RF. 2004. Pressure inactivation of *Bacillus* endospores. Appl Environ Microbiol 70:7321–7328. 10.1128/AEM.70.12.7321-7328.200

63 Ning Z, Dong W, Bian Z, Huang H, Hong K. 2024. Insight into effects of terbium on cell growth, sporulation and spore properties of *Bacillus subtilis*. World J Microbiol Biotechnol 40:79. 10.1007/s11274-024-03904-4

64 Hrustić J, Mihajlović M, Stevanović M, Gašić S, Grahovac M, Tanović B. 2023. Effectiveness of an indigenous *Bacillus subtilis* B6 strain in the control of postharvest apple fruit rot. Eur J Plant Pathol 167:727–742. 10.1007/s10658-023-02719-7

65 Restuccia C, Giusino F, Licciardello F, Randazzo C, Caggia C, Muratore G. 2006. Biological control of peach fungal pathogens by commercial products and indigenous yeasts. J Food Prot 69:2465–2470. 10.4315/0362-028x-69.10.2465

66 Wang SY, Herrera-Balandrano DD, Wang YX, Shi XC, Chen X, Jin Y, Liu FQ, Laborda P. 2022. Biocontrol ability of the *Bacillus amyloliquefaciens* group, *B. amyloliquefaciens*, *B. velezensis*, B. nakamurai, and B. siamensis, for the management of fungal postharvest diseases: a review. J Agric Food Chem 70:6591–6616. 10.1021/acs.jafc.2c01745

67 Wang B, Wang S, Geng QR, Zhang NH, Zhuo QH, Zhou QR, Zeng H, Tian J. 2024. Effects of perillaldehyde and polyamines on defense mechanisms of sweet potatoes against *Ceratocystis fimbriata*. J Agric Food Chem in press. 10.1021/acs.jafc.4c07055

68 Deng HY, Wang FR, Wu Q, Sun HJ, Ma JK, Ni R, Li ZY, Zhang LM, Zhang J, Liu MY. 2024. Novel multiresistant osmotin-like protein from sweetpotato as a promising biofungicide to control *Ceratocystis fimbriata* by destroying spores through accumulation of reactive oxygen species. J Agric Food Chem 72:1487– 1499. 10.1021/acs.jafc.3c07663

69 Liu MY, Gong Y, Sun HJ, Zhang J, Zhang LM, Sun J, Han YH, Huang JJ, Wu Q, Zhang CL, Li ZY. 2020. Characterization of a novel chitinase from sweet potato and its fungicidal effect against *Ceratocystis fimbriata*. J Agric Food Chem 68:7591–7600. 10.1021/acs.jafc.0c01813

